# Wilting Wildflowers and Bummed-Out Bees: Climate Change Threatens U.S. State Symbols

**DOI:** 10.1101/2024.09.08.611901

**Authors:** Xuezhen Ge, Ya Zou, Heather A. Hager, Jonathan A. Newman

## Abstract

Species designated as state symbols in the United States carry cultural importance and embody historical heritage. However, they are threatened by climate change and even face the risk of local or global extinction. The responses of these species to climate change have received little attention. In this study, we examine the effects of climate change on 64 state flowers and 68 state insects in the United States by employing correlative species distribution models (SDMs). We select a variety of commonly used SDM algorithms to construct an ensemble forecasting framework aimed at predicting the potential climatic habitats for each species under both historical (1981-2010) and future (2071-2100) climate scenarios (SSP1-2.6 and SSP5-8.5), and how these changes might influence the habitat suitability of flower and insect species within their symbolic states and across the United States. Our results indicate that 30 − 66% of state flowers and 18 − 51% of state insects are projected to experience substantial losses of climatically suitable habitat within their symbolic states. Under the high-emissions scenario (SSP5-8.5), ten state flowers and three state insects are likely to face local extinction by the 2080s. Although most of these species may find suitable habitats in other states, only two are projected to have such areas located adjacent to their current symbolic states, potentially limiting natural dispersal. Nationally, 85% of flower species and 71 − 79% of insect species are expected to shift their suitable habitat both poleward and uphill, with the magnitude of latitudinal and elevational shifts significantly greater under SSP5-8.5 than under SSP1-2.6. These findings highlight the vulnerability of culturally significant species to climate change and underscore the urgency of integrating climate adaptation into conservation planning. Proactive, forward-looking conservation and management strategies may be critical for preserving cultural heritage and maintaining ecosystem resilience.

## INTRODUCTION

American states have official flowers and insects [and often other flora and fauna; 1, 2, 3]. Some states share the same official species, while others boast unique species exclusive to their region. Writing about state flowers and trees, Nord [1] says:

> “The history behind our nation’s selection of its flowers and trees is rich with political intrigues, legends, deception, and humor, which makes each state’s adoption a unique story.” (p. xiii)

Presumably, the states feel these species are in some way particularly representative of their citizens’ way of life—perhaps they are common (e.g., peach blossom *Prunus persica* in Delaware) or once were and are now endangered (e.g., showy lady’s slipper *Cypripedium reginae* in Minnesota, rusty patched bumble bee *Bombus affinis* in Minnesota), or perhaps the majority of citizens just have a fondness for these species (e.g., black-eyed Susan *Rudbeckia hirta* in Maryland). Why they were chosen differs by state, but these species are in some way *culturally significant* to citizens of the state. Jackson and Perkins [4] write:

> “The diverse array of state flowers in the United States weaves a tapestry of natural beauty, cultural significance, and historical heritage. These floral emblems encapsulate the essence of each state, reflecting its unique landscapes, economic contributions, and regional identity.”

These unique histories and contributions are threatened when state flora and fauna are endangered by climate change. Developed nations such as the United States often spend large sums to conserve their cultural and historical heritage. Research on climate change is not new, but the impact of climate change on cultural heritage has only recently emerged as a subject of research.

This issue gained attention at the Conference of the Parties to the United Nations Convention on Climate Change in Madrid in 2019 (COP25) and continues to be a prominent topic of interest [5, 6]. Most commonly, researchers have considered *cultural heritage* in the form of the so-called “tangible cultural heritage”, such as architecture or ancient anthropological sites. Nevertheless, intangible cultural heritage, including customs and Indigenous knowledge, also faces threats just as significant due to climate change [7].

Climate change is driving geographical shifts in species distributions, with many organisms migrating northward in latitude or upward in elevation to remain within their preferred climate niche [8, 9]. Although biogeographic responses to climate warming have been widely documented across diverse taxa, the range shifts of culturally significant species—and their potential ecological and sociocultural impacts—remain largely understudied. As climate conditions become progressively unfavorable in their traditional habitats, certain culturally significant species might become extirpated in the states they presently symbolize and/or relocate to other states with more suitable conditions. These shifts can disrupt the cultural bond between species and local customs, affect long-established educational and cultural traditions, and weaken the public’s connection to their natural heritage [10]. Moving to new states, these species may encounter inadequate protections and management issues due to current conservation policies being designed around historical, rather than future, ranges [11]. Such range shifts may also lead to new ecological interactions, habitat fragmentation, localized population declines, and increased ecosystem vulnerability [12]. Identifying climate change impacts on these culturally important species is crucial for future cultural heritage preservation and maintaining the symbolic connections between communities and their natural environments.

In this study, we employ various commonly used correlative species distribution model (SDM) algorithms with an ensemble forecasting framework to predict potential distributions of 64 state flowers and 68 state insects under historical and future climate scenarios. It is important to note that some states have both “state flowers” and “state wildflowers”, while others may designate more than one flower or insect. For a complete list of species, see Supplementary Tables S1–S3. By modeling changes in habitat suitability across the United States, we aim to address three key questions: (1) Will these symbols of cultural heritage—officially adopted by their states—continue to persist within state boundaries and uphold their multifaceted symbolic roles? (2) Are these species likely to shift beyond their symbolic states (i.e., the states where they serve as official state species), potentially weakening symbolic ties and underscoring the need for coordinated conservation strategies? (3) Do these species exhibit consistent directional shifts—particularly northward or uphill—that reflect known biogeographic responses to climate warming and highlight emerging conservation priorities? Our study contributes not only to informing proactive conservation and cultural adaptation strategies, but also to identifying whether alternative states may emerge as suitable refuges to support the continued survival and symbolic relevance of these species.

## MATERIALS AND METHODS

### Species lists

We compiled lists of official state flower and insect species by consulting netstate.com and cross-referencing with associated state legislation and state websites (See Supplement 1 and 2). Flowers included 46 state flowers, 15 state wildflowers, 1 state floral emblem, 1 children’s state flower, and 1 beautification and conservation plant (total = 64; ∼ 83% native species; Supplementary Table S1). Four state flowers were excluded due to the lack of occurrence records: Hawaii’s endemic endangered *Hibiscus brackenridgei*, New York’s cultivated *Rosa* spp., Ohio’s cultivated *Dianthus caryophyllus* (carnation), and Oklahoma’s cultivated hybrid tea rose ‘Oklahoma’ (*Rosa* × *hybrida*). Of the 64 official flowers included, 9 were designated by genus alone or as multiple species (Florida, *Coreopsis* spp.; Georgia, native *Rhododendron* spp.; Illinois, *Asclepias* spp.; Indiana, cultivated *Paeonia* spp.; Michigan, *Malus coronaria* and *Malus domestica/pumila*; Mississippi, *Coreopsis* spp.; New Mexico, *Yucca* spp.; North Dakota, *Rosa arkansana* and *Rosa blanda*; Texas, *Lupinus* spp.). For genera, we determined all the species native to the state using the USDA Plants database and included those with sufficient occurrence records for modelling (Supplementary Table S2). For cultivated *Paeonia*, we included *P. lactiflora* and *P. officinalis*. After accounting for all species selected for states that did not designate a specific species name, we included 114 floral species and modelled their habitat suitability under climate change.

Insects included 41 state insects, 21 state butterflies, 3 state agricultural insects, 1 state bee, 1 state bug, and 1 state pollinator (total = 68; ∼ 68% native species; Supplementary Table S3). Another state bug was excluded (Delaware, *Coccinella* spp.) because the genus was too broad. When only a common name or genus was specified in the legislation, we consulted other state materials to determine if there was an accepted representative species. Of the 68 insect species analyzed, many species are designated by multiple states. For example, European honey bee (*Apis mellifera*) is recognized by 20 states, and the monarch butterfly (*Danaus plexippus*) by 7 states. After accounting for the multiple uses of these species, we included 34 insect species and modelled their habitat suitability under climate change.

### Species occurrence records

We used the `R` package, `rgbif` [13], to obtain global occurrence records for all species from the Global Biodiversity Information Facility (GBIF, http://www.gbif.org), accessed December 2023. Following the methods of Ge et al. [14, p4], we removed duplicate records and those with spatial and temporal errors using the `R` package, `CoordinateCleaner`[15]. We spatially thinned the raw occurrence data by randomly selecting a single presence point within each 10 × 10 km grid cell using the `R` package, `BiodiversityR`, to reduce sampling bias [16] (See Supplementary Tables S1-S3 for the number of occurrence records used during data cleaning and thinning.). Depending on the correlative model algorithm and the number of occurrence records [17], we used different strategies [Supplementary Figure S1, 14] to generate the pseudo-absence data for each species using the `R` package, `biomod2` [18]. Finally, we split the occurrence and pseudo-absence data sets into training sets for model fitting and test sets for model evaluation using the block cross-validation technique, implemented in the `R` package, `blockCV` [19]. The processes used to generate pseudo-absence data and split the training and test data sets are listed in Supplementary Figure S2 of Ge et al.’s study [14].

### Climate data

We downloaded 19 raster-based bioclimate variables from CHELSA V2.1 (Climatologies at High Resolution for the Earth’s Land Surface Areas, https://chelsa-climate.org/, [20]) to represent the historical (1981–2010) and future (2071–2100) climates. The historical climate data included in CHELSA V2.1 were generated by downscaling temperature and precipitation estimates from the ERA-Interim climatic reanalysis to a high resolution of 30 arcseconds [20]. CHELSA V2.1 also includes future bioclimate variables that were derived for five global climate models (GCMs: GFDL-ESM4, IPSL-CM6A-LR, MPI-ESM1-2-HR, MRI-ESM2-0, and `UKESM1-0-LL`) and three Shared Socioeconomic Pathways (SSPs: SSP1-2.6, SSP3-7.0, and SSP5-8.5) as well as three time periods (2011–2040, 2041–2070, 2071–2100). We selected the two scenarios (SSP1-2.6 and SSP5-8.5), as we are interested in the range of climate change effects under different degrees of warming. Historical (1981–2010) and future (2071–2100) global climate data are used here to demonstrate how species’ habitat suitability changes over time. The multi-model ensemble can reduce the uncertainty that results from differences in the GCMs [21, 22], so we calculated the ensemble mean value of the five GCMs and used these values to generate future global projections with the SDMs. Given the global study area, we ‘upscaled’ each bioclimatic variable from 1 km to 10 km resolution. For the directional shift analysis described below, we obtain the elevation data from AWS Terrain Tiles (derived from NASA’s Shuttle Radar Topography Mission) via `elevatr` package at zoom level 5, and then resample projected species distribution rasters using bilinear interpolation to match the resolution of this elevation raster.

### Ensemble correlative Species Distribution Model (SDM)

Among numerous correlative SDM algorithms, we selected eight that are commonly used and provided by the `Biomod2` package: generalized linear models [GLM, 23], generalized additive models [GAM, 24], generalized boosted models [GBM, 25], multivariate adaptive regression splines [MARS, 26], classification tree analysis [CTA, 27], flexible discriminant analysis [FDA, 28], and maximum entropy [MaxEnt, 29].

Following the methods of Ge et al. [14, p4], we applied model-specific training and test data sets for presence and pseudo-absence data across various model algorithms for each species. For each presence training set and its corresponding historical climate data, we used the variance inflation factor (VIF) to assess the multicollinearity between the 19 bioclimate variables and the excluded variables with VIF > 4. The remaining variables were used to fit the model to historical data, and the fitted model, combined with future climate values, predicted future climatic habitat ranges. We generated multiple training and test data sets for each algorithm, with each replicate representing a model simulation. Simulations were evaluated using a 10% omission rate, excluding those where more than 10% of the test data appeared in unsuitable climatic habitat to prevent overfitting. We then averaged habitat suitability across all locations for both historical and future conditions. For future predictions (2071–2100, SSP1-2.6 and SSP5-8.5), we used validated models that met the omission rate criteria. The model projections for each state species are included in SI Dataset (https://osf.io/j8qnh/?view_only=dde6f42192b14339ac8fb38b604609d5).

### Measuring climate change effects on habitat suitability within symbolic states

We used the ensemble correlative SDM approach described above to predict habitat suitability values (HSV) of species within the United States under historical (1981–2010) and future (2071– 2100) climate conditions. For the states that only designated a genus, we selected species with sufficient occurrence records for modelling from all the species native to the state using the USDA Plants database, and then determined the HSVs of the genus by calculating the maximum HSV among all the selected species ([1, …, *n*]) in each grid cell (*i*) using:

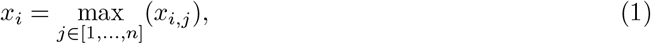

where *x*_*i*_ represents the habitat suitability value for the state species in grid cell *i*. We did the same calculation for all states with multiple species.

For the 64 state flowers and 68 state insects, we obtained the habitat suitability value of each grid cell over two time periods and two SSP scenarios within their respective states and compared the distributions of their habitat suitability values under climate change. Furthermore, we compared the effects of climate change on the habitat suitability of species within their symbolic states using the following three metrics:

#### 1. Change in median (δ)

We applied the bootstrap method to estimate the difference in median values and their 95% confidence intervals. The 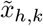 and 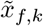 are the medians of the *k*^*th*^ (*k* ∈ [1, …, 10_4_], repeat 10_4_ times) bootstrap sample of all habitat suitability values under historical (*h*) and future (*f*) climate conditions, respectively. The *δ*_*k*_ is the difference in the median of the *k*^*th*^ bootstrap sample under climate change; *δ* is the bootstrap estimate of the median difference between the two climate conditions.

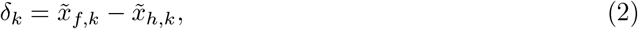

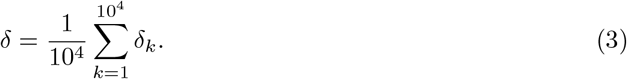

Then,

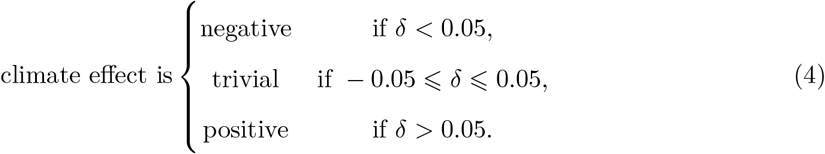

#### 2. Spatial variation (*CV*)

The coefficient of variation (*CV*) can quantify the extent of spatial variability in habitat suitability across the state. We estimated *CV* for each state species by calculating the difference in habtat suitability for each grid cell (Δ*x*_*i*_, Equation 5), and the mean (*µ*, Equation 6) and standard deviation (*σ*, Equation 7) of these difference values. *CV* < 1 indicates small spatial variation, whereas *CV* ⩾ 1 denotes large spatial variation.

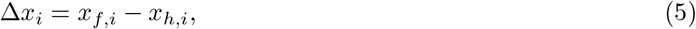

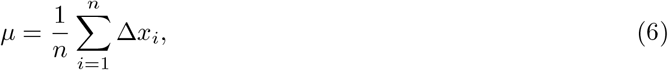

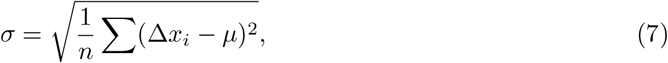

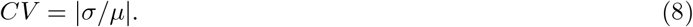

#### 3. Local extinction indicator (*θ*)

We define the local extinction indicator (*θ*) based on the median in habitat suitability values under future climate condition 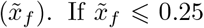, we classify the state species as likely to face local extinction in this state.

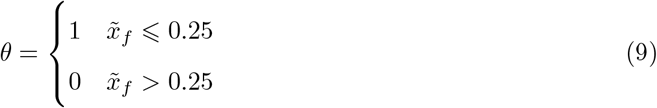

Based on the values of *δ, CV* , and *θ*, we categorized the responses of the state species to climate change into six distinct types:

- **Negative** overall effect causes local *extinction*.
- **Negative** overall effect *reduces* suitability.
- **Trivial** overall effect with *small* spatial variation.
- **Trivial** overall effect with *large* spatial variation.
- **Positive** overall effect with *small* spatial variation.
- **Positive** overall effect with *large* spatial variation.

### Identify alternative suitable states for designated species

To identify alternative suitable states for species, we first identify symbolic states where the projected median future habitat suitability value 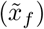 falls below 0.25, indicating a high risk of local extinction under future climate scenarios (Local extinction indicator *θ* = 1, Equation 9). For these species, we then calculate the value of 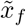 in all other states to identify potential new suitable climatic habitats. States where the projected median habitat suitability 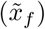 under the high emissions scenario (SSP5-8.5) exceeds 0.5 are considered alternative suitable states, and their count is denoted as *N*_alt_. To assess potential gains, we compare the median habitat suitability values under historical and future climates across all states and count those where suitability increases (*δ* > 0, Equation 3) under climate change, denoted as *N*_gain_.

### Assessing spatial changes in habitat suitability with the U.S

#### 1. Directional shift analysis

We assess latitudinal and elevational shifts in areas of high habitat suitability by comparing the centroid of suitable grid cells under historical and future climate conditions. Specifically, we extract grid cells where the habitat suitability value (*x*_*i*_) exceeds 0.5 in both time periods. For each time period *j* ∈ {*h, f*}, we define the set of highly suitable grid cells as:

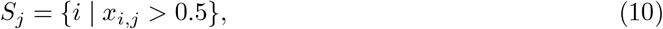

where *x*_*i,j*_ is the habitat suitability value at grid cell i under historical (*j* = *h*) or future (*j* = *f*) climate. For each period, we then calculate the weighted mean latitude 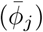 and elevation 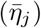 of these suitable cells to represent the spatial centroid of highly suitable climatic habitat:

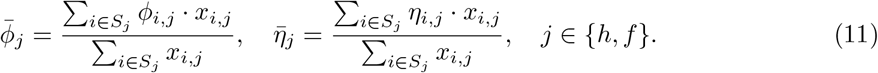

Here, *ϕ*_*i,j*_ and *η*_*i,j*_ are the latitude and elevation of grid cell *i* under period *j*. The latitudinal and elevational shifts under climate change are then quantified as:

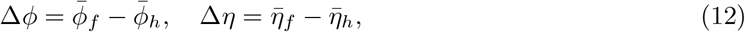

where Δ*ϕ* > 0 indicates a northward shift and Δ*η* > 0 indicates an uphill shift.

#### 2. Net change (gain-loss) analysis

Most SDM studies estimate the net change in species’ potential distributions under climate change by evaluating how climatic habitat is projected to expand, contract, or remain stable over time [30, 31]. This is typically done by converting continuous habitat suitability values from historical and future projections into binary maps (suitable vs. unsuitable) using predefined thresholds, such as the maximum training sensitivity plus specificity or the 10th percentile training presence [32, 33]. From these binary maps, three components are commonly derived: climatic habitat gain (areas projected to become suitable in the future but were unsuitable historically), climatic habitat loss (areas projected to become unsuitable in the future but were suitable historically), and stable climatic habitat (areas suitable in both time periods).

However, in our study, we avoid discretizing habitat suitability because applying the same threshold across all species would ignore ecological differences, while using species-specific thresholds would limit comparability across species. Instead, we directly calculate the difference in habitat suitability between future and historical projections for grid cell *i* as Δ*x*_*i*_ = *x*_*i,f*_ − *x*_*i,h*_ [34]. We define climatic habitats as stable if |Δ*x*_*i*_| ≤ 0.01, representing minimal change given our rounding precision (two decimal places). Grid cells with Δ*x*_*i*_ > 0.01 are classified as climatic habitat gain, while those with Δ*x*_*i*_ < −0.01 are classified as climatic habitat loss. Net change is then quantified as the difference between the proportion of habitat gain and the proportion of habitat loss for each species.

## RESULTS

### Habitat loss and extinction risk within symbolic state ranges

Figures 1A–B and 2A–B illustrate projected changes in median habitat suitability values and their distributions for state flowers and insects under climate change. The majority of these state species are expected to experience declining habitat suitability over time, increasing the risk of local extinctions by the 2080s (Table 1, Figures 1A and 2A). Under the high emissions scenario, which assumes CO_2_ emissions triple by 2075 (SSP5-8.5), 42 (∼ 66%) state flowers and 35 (∼ 51%) state insects will face reduced habitat suitability, with the median habitat suitability value decreasing by up to 83%, e.g., showy lady’s slipper (*C. reginae*) in Minnesota. In total, ten (∼ 16%) state flowers and three (∼ 4%) state insects are projected to lose so much climatically suitable habitat under this scenario, that the losses suggest the potential local extinction (Table 2).

**Table 1:**
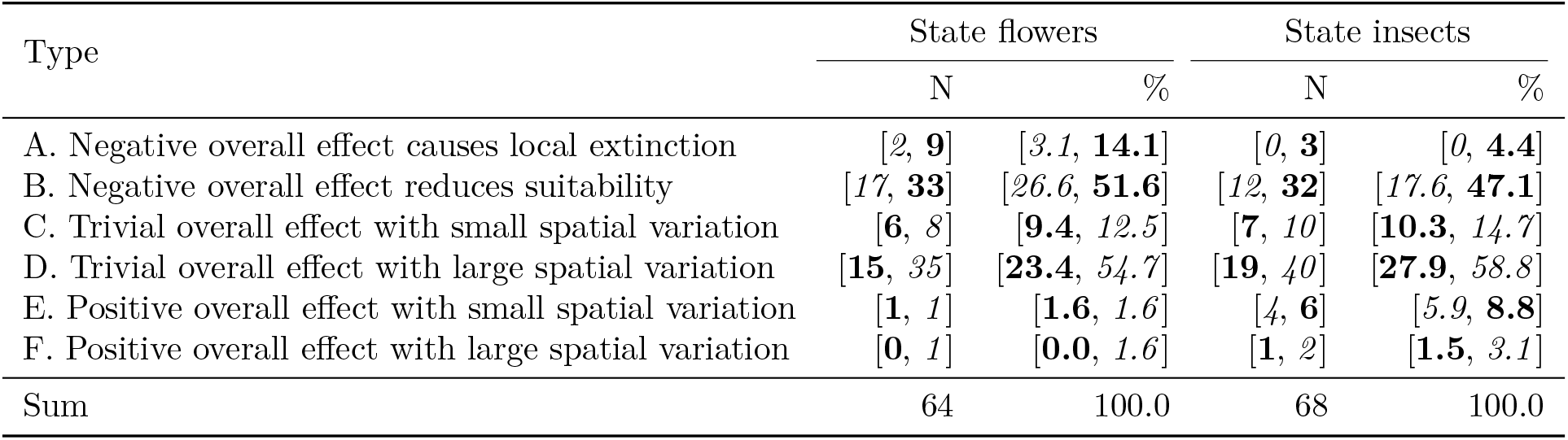
Quantities and proportions of state flowers and state insects across various types of responses to climate change (see Methods for classification criteria). The numbers in italic and bold represent predicted results under SSP1-2.6 and SSP5-8.5, respectively. Some states designate only a species group rather than a specific species; in these cases, we treat multiple species as a single entity by selecting the one with the highest habitat suitability within each grid cell. Additionally, some states share the same species as their state symbol. In total, the analysis includes 64 state flowers and 68 state insects.

**Table 2:**
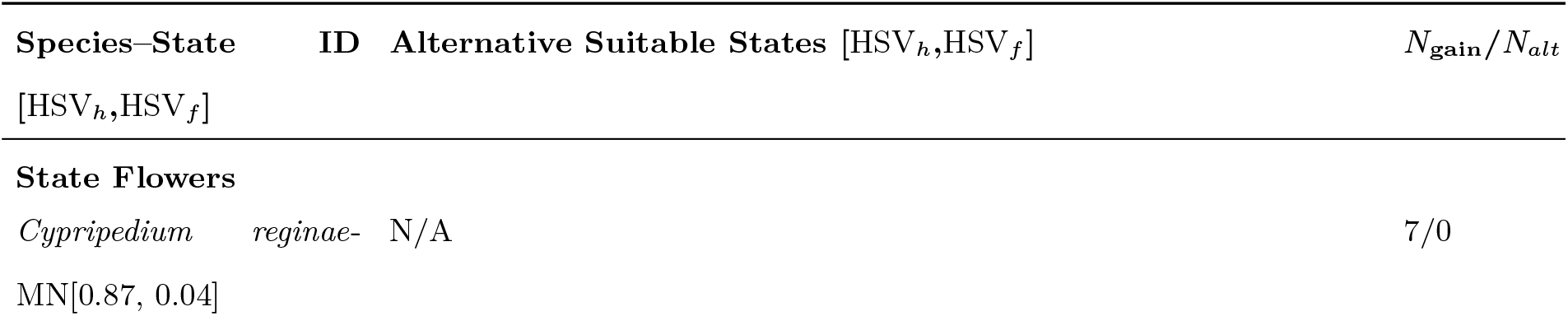
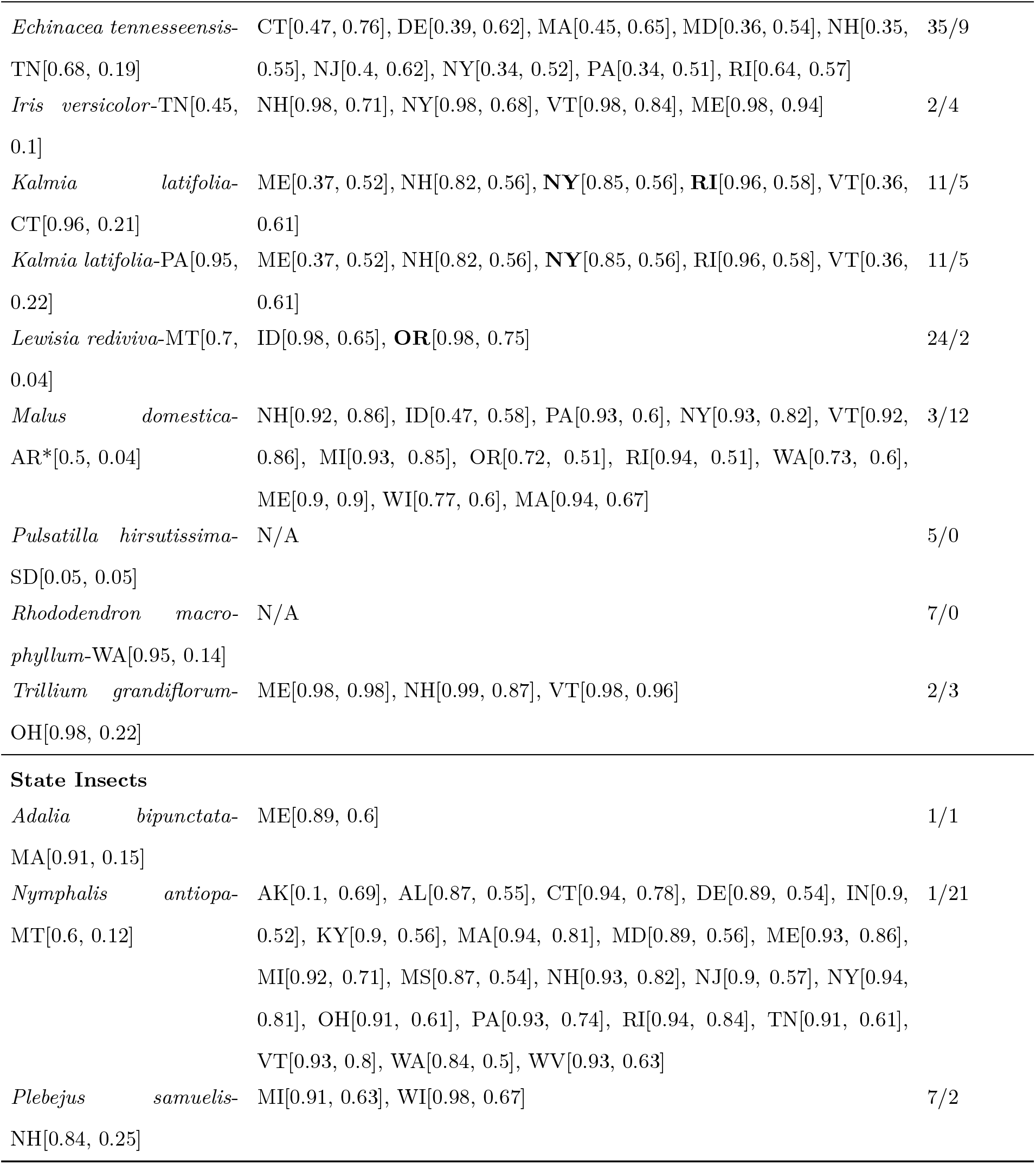
Alternative suitable states for selected state flowers and insects under the SSP5-8.5 scenario by 2100. *Species-State* column lists the state species whose median habitat suitability values (HSVs) are below 0.25 within their symbolic states by the 2080s. *Alternative suitable states* represent the non-symbolic states whose median HSV_*f*_ > 0.5. *N*_alt_ denotes the number of such alternative suitable states, and *N*_gain_ indicates the number of states where median suitability increases under climate change. *N/A* indicates no new suitable states identified. Among the alternative suitable states, those that are geographically adjacent to the symbolic state are shown in bold.

**Figure 1:**
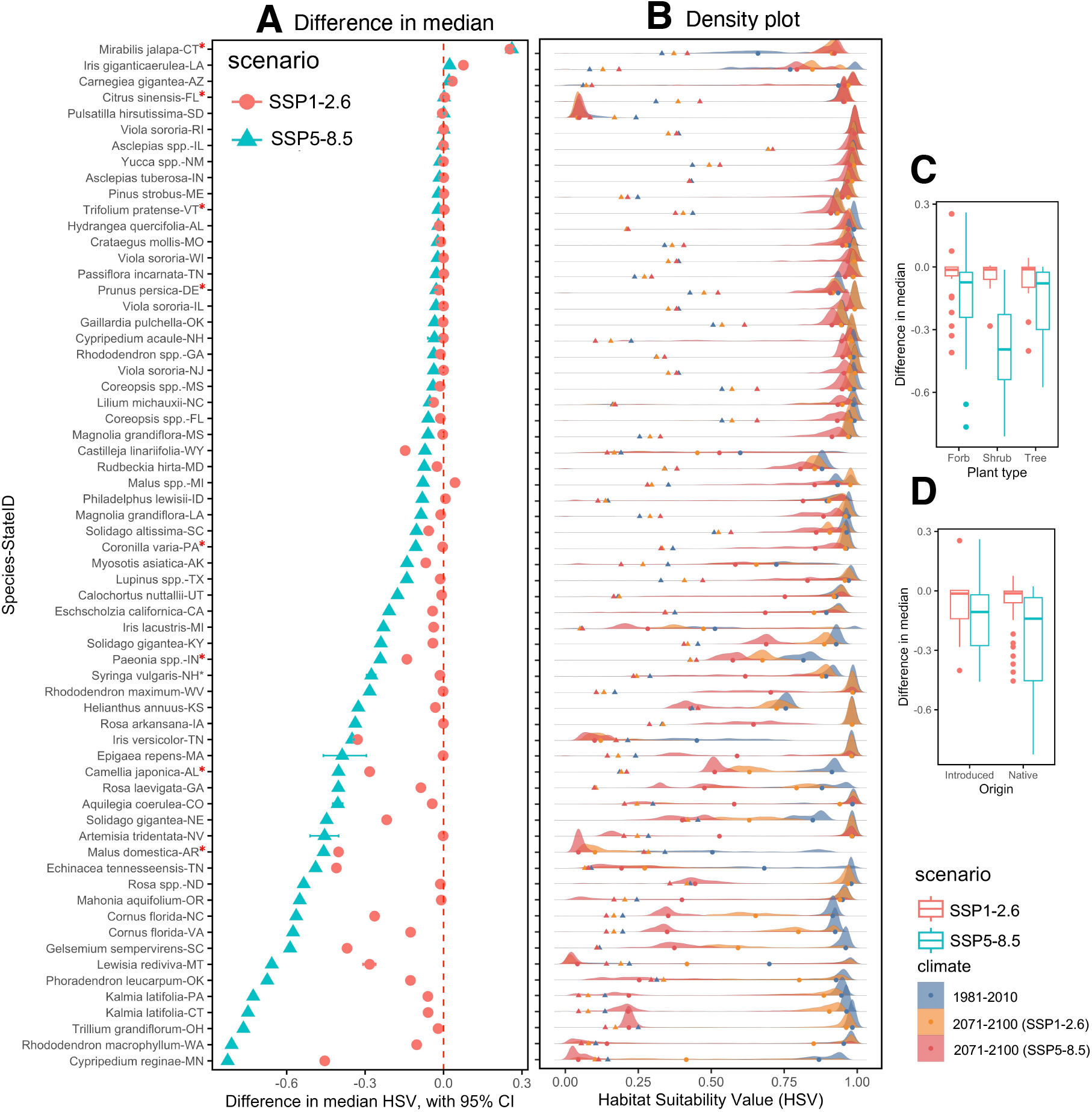
State flowers: Changes in habitat suitability for 64 state flowers under climate change. Panel A shows the difference in the median habitat suitability value (HSV) between future and historical climate conditions under two SSP scenarios (SSP1-2.6 and SSP5-8.5) for each state flower in its symbolic state. These state species are ranked by the difference value under SSP5-8.5 scenario. Non-native species are indicated by red ⋆ in y-axis labels. Panel B plots the distribution of HSVs under different time periods and different SSP scenarios (see figure legends); colored circular dots represent the median HSVs within the symbolic state, while colored triangular dots indicate the mean HSVs across all grid cells within the United States. Panels C-D show box plots of difference values for all state flowers grouped by plant type (C), origin (D). Large differences in sample size could affect the reliability of comparisons.

Even under the low emissions scenario, which assumes CO_2_ emissions cut to net zero around 2075 (SSP1-2.6), 19 (∼ 30%) state flowers and 12 (∼ 18%) state insects would experience negative climate effects. Although, *on average*, a large portion of state species (state flowers: 32 − 69%; state insects: 38 − 74%) may be largely unaffected by climate change, these effects vary geographically within individual states (Table 1, Figure S1 and Figure S2), with approximately 23 (∼ 55%) flowers and 28 (∼ 59%) insects expected to exhibit considerable geographical movement caused by climate change. Some, such as the giant blue iris (*Iris giganticaerulea*), show more pronounced and obvious spatial variations in their response to change than others (Figure S1D).

We also examined whether climate change impacts different groups of species unevenly, focusing on the SSP5-8.5 scenario due to its more pronounced effects. Among state flowers, shrubs are generally more negatively affected than forbs or trees (Figure 1C), and native species are more vulnerable than introduced species (Figure 1D). For state insects, butterflies experience slightly greater negative impacts than bees or ladybugs (Figure 2C), and—as with flowers—native insects are more susceptible to climate change than non-native species (Figure 2D).

**Figure 2:**
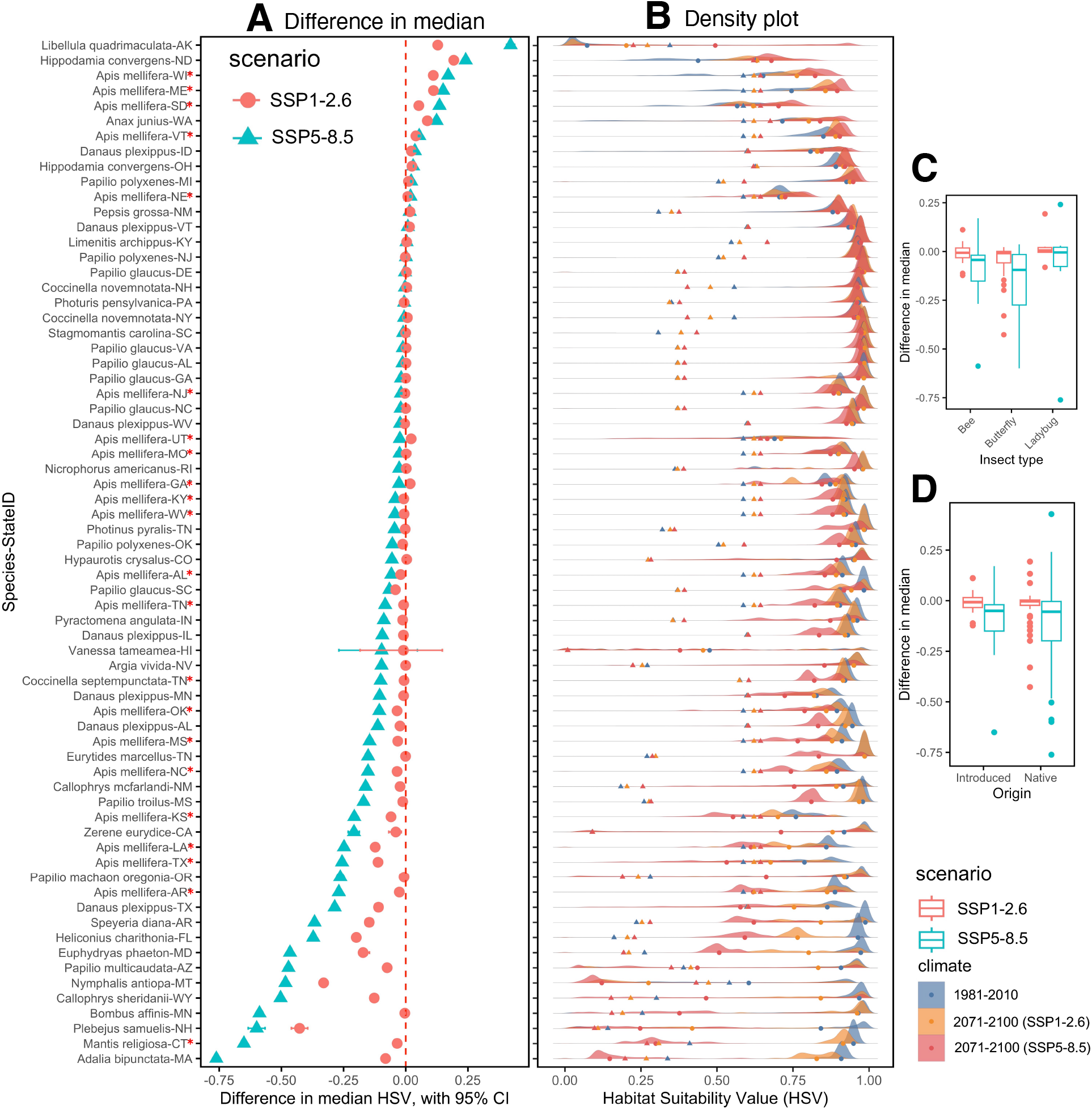
State insects: Changes in habitat suitability for 68 state insects under climate change. Panel A shows the difference in the median habitat suitability value (HSV) between future and historical climate conditions under two SSP scenarios (SSP1-2.6 and SSP5-8.5) for each state insect in its symbolic state. These state species are ranked by the difference value under SSP5-8.5 scenario. Non-native species are indicated by red ⋆ in y-axis labels. Panel B plots the distribution of HSVs under different time periods and different SSP scenarios (see figure legends); colored circular dots represent the median HSVs within the symbolic state, while colored triangular dots indicate the mean HSVs across all grid cells within the United States. Panels C-D show box plots of difference values for all state insects grouped by insect type (C), origin (D). Large differences in sample size could affect the reliability of comparisons.

### Emerging suitable states under climate change

Among the ten state flowers and three state insects projected to face local extinction due to climate change, we identify alternative states where the median HSV is projected to remain above 0.5 by the 2080s. These are considered newly emerging suitable climatic habitats and are listed in Table 2. Notably, three state flowers—the showy lady’s slipper (*C. reginae*), the pasque flower (*Pulsatilla hirsutissima*), and the rhododendron (*Rhododendron macrophyllum*)—are projected to have no suitable climatic habitat anywhere in the United States by the 2080s. Historically, the showy lady’s slipper and the pasque flower had relatively broad suitable ranges, with 13 and 20 states respectively showing median HSVs above 0.5 (Supplementary Table S4). In contrast, the rhododendron had a more restricted range, with only five states exceeding the 0.5 threshold historically and none projected to do so under the SSP5-8.5 scenario by the 2080s.

Although some state species have a variety of alternative suitable climatic habitats for colonization, such as Arkansas’s state flower, the domesticated apple (*Malus domestica*), which has 12 other viable states, and Montana’s state insect, the mourning cloak (*Nymphalis antiopa*), with 21 potential states, just three state flowers have suitable areas that are geographically adjacent to their symbolic states (Table 2, Figure 3F4-6). The other seven state flowers and three state insects lack nearby suitable environments (Figure 3). Among the 13 state species, states showing an increase in median HSV due to climate change are significantly fewer compared to those showing a decrease (N_gain_ ≪ 25). This suggests that many state species may face limited access to climatically suitable regions once their symbolic states become unsuitable. Our findings highlight the severity of projected climatic habitat loss for these species under future climate scenarios.

**Figure 3:**
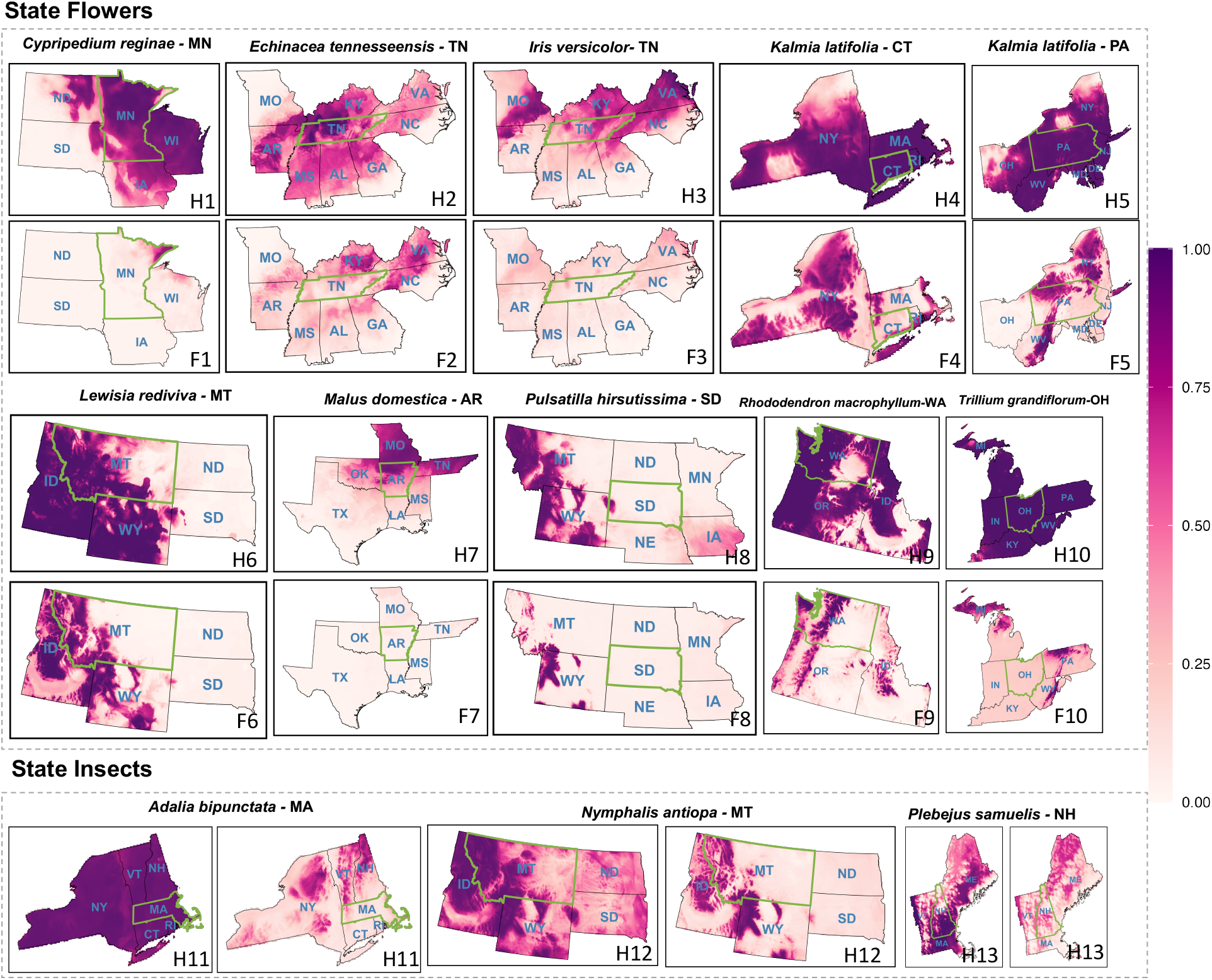
Habitat suitability maps for ten state flowers and three state insects (listed in Table 2) under historical and future climate conditions (SSP5-8.5 scenario). Each panel shows the spatial distribution of habitat suitability within a species’ symbolic state and its adjacent states. Darker colors indicate higher suitability. In plot labels, ‘H’ refers to historical climate (1981–2010), and ‘F’ refers to future climate projections under the SSP5-8.5 scenario (2071–2100).

### Northward and uphill: consistent biogeographic responses to warming

Among the 114 individual flower species and 34 individual insect species included in this study, changes in mean habitat suitability (HSV) show largely consistent directional trends across climate scenarios. Specifically, 45 (∼ 39%) flower species and 17 (∼ 50%) insect species are projected to experience increased mean HSV under both SSP1-2.6 and SSP5-8.5 scenarios, while 53 (∼ 46%) flower species and 11 (∼ 32%) insect species are projected to experience decreases under both scenarios (Figure 4A, Supplementary Table S5). The SSP5-8.5 scenario results in a substantially stronger impact on mean HSV—approximately twice the magnitude observed under SSP1-2.6 (Figure 4A). Directional shift analysis reveals clear northward and uphill migration patterns among these culturally significant species (Figure 4B-C). Under the SSP1-2.6 scenario, the centroid of highly suitable climatic habitat shifts to higher latitudes for 111 (∼ 97%) flower species and 33 (∼ 97%) insect species, and to higher elevations for 100 (∼ 88%) flower species and 28 (∼ 82%) insect species. Under the SSP5-8.5 scenario, the latitudinal centroid shifts northward for 112 (∼ 98%) flower species and 33 (∼ 97%) insect species, while 99 (∼ 87%) flower species and 24 (∼ 71%) insect species show upward elevational shifts. In total, 85% of flower species and 74 − 79% of insect species are projected to shift both poleward and uphill under two emission scenarios (Supplementary Table S6). While the proportion of shifting species is comparable between scenarios, the *magnitude* of both latitudinal and elevational shifts is significantly greater under SSP5-8.5. These results highlight strong and consistent poleward and elevational responses to climate change, with more pronounced shifts under the high-emissions scenario. When comparing flowers and insects, flower species tend to shift farther northward and to higher elevations than insect species. For example, the elevational centroid of highly suitable climatic habitat for the pasque flower is projected to rise from 1,150 m to 2,279 m under SSP5-8.5 by the 2080s. Similarly, the latitudinal centroid for the showy lady’s slipper is expected to move from 43.8°N to 63.2°N, indicating a dramatic poleward shift.

**Figure 4:**
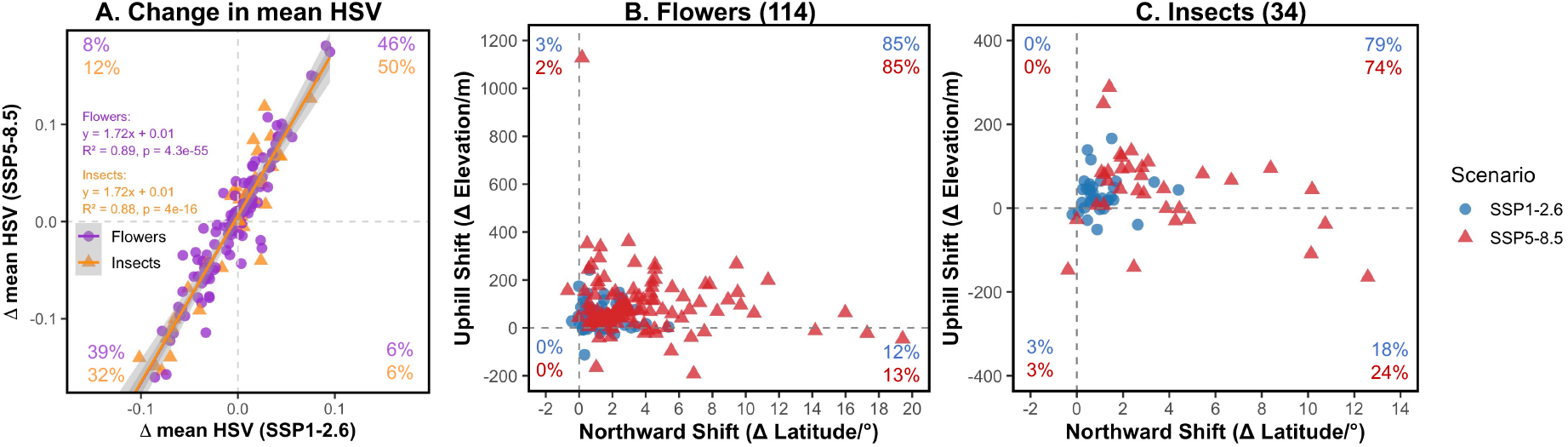
Climate-driven shifts in habitat suitability and distribution patterns for 114 flower species and 34 insect species under SSP1-2.6 and SSP5-8.5 scenarios. (A) Change in mean habitat suitability value (HSV) between historical and future climate for state flowers (purple circles) and state insects (orange triangles). Linear regression lines and equations are shown for each group. Percentages in each quadrant indicate the proportion of species showing consistent increases (top right), consistent decreases (bottom left), or mixed responses in mean HSV across scenarios. (B, C) Latitudinal and elevational shifts in the centroid of highly suitable habitat for flowers (B) and insects (C) under SSP1-2.6 (blue circles) and SSP5-8.5 (red triangles). Percentages in each quadrant reflect the proportion of species showing directional trends in shift patterns.

We also conduct a net habitat change analysis by comparing the projections under historical climate and future climate with SSP5-8.5 scenario, visualized using a circular bar plot that illustrates the difference between climatic habitat gain and loss for each species (Figure 5). Different colors distinguish species groups, while pairs of adjacent bars with varying color intensities represent net change within the symbolic state and across the entire United States, respectively. Nationally, approximately 63% of state flowers and 31% of state insects are projected to experience greater habitat loss than gain. Within their symbolic states, this pattern is even more pronounced: about 93% of state flowers and 76% of state insects are expected to lose more climatic habitat than they gain (Supplementary Table S7). This pattern remains consistent across taxa, as species-rich groups also show a majority of species experiencing net habitat loss under future climate scenarios. Moreover, the magnitude of net change is typically greater within symbolic states compared to the entire United States, highlighting the heightened vulnerability of species within their designated cultural ranges.

**Figure 5:**
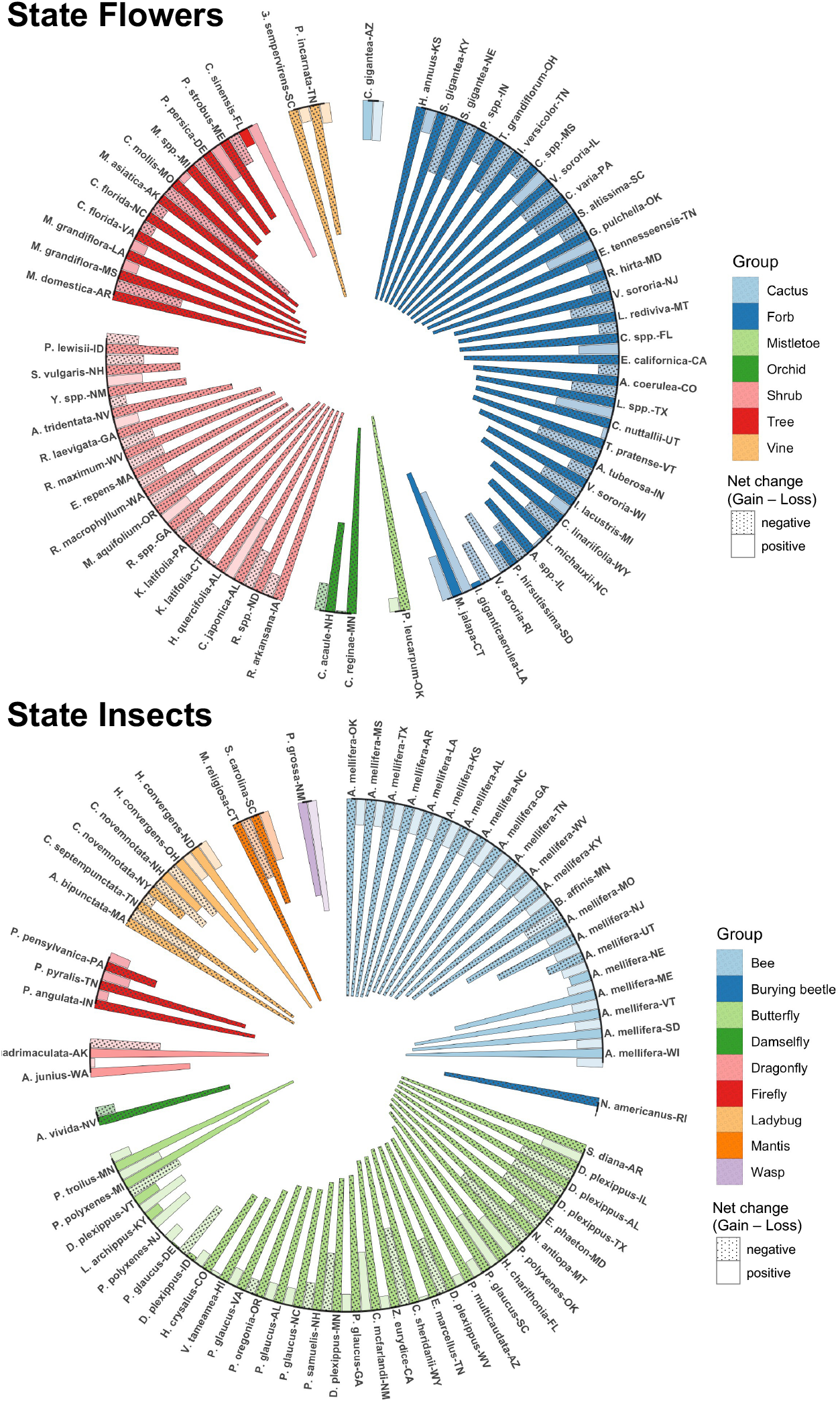
Net climatic habitat change for state flowers (top) and state insects (bottom) under the SSP5-8.5 climate scenario. Species are grouped by taxonomic category (e.g., Tree, Forb, Bee, Butterfly), indicated by bar color. Each symbolic species is represented by two bars: a darker bar indicating net change within its symbolic state, and a lighter bar representing net change across the entire United States. Bar length corresponds to the net climatic habitat change, defined as the difference between the proportion of habitat gain and habitat loss projected by 2100 relative to the historical baseline (1981–2010). Instead of converting continuous habitat suitability values into binary suitable/unsuitable maps using arbitrary thresholds, we quantified change directly as per-pixel suitability differences (Δ*x*_*i*_ = *x*_*i,f*_ −x_*i,h*_). Grid cells with |Δ*x*_*i*_| ≤ 0.01 were classified as stable, while cells with Δ*x*_*i*_ > 0.01 or Δ*x*_*i*_ < − 0.01 were categorized as habitat gain or loss, respectively. Bars with a dotted fill pattern indicate species for which the projected loss of suitable habitat exceeds the projected gain, resulting in a negative net change.

## DISCUSSION

Our study examined the impacts of climate change on species of cultural significance—specifically, United States state flowers and insects. We identified ten state flowers and three state insects that will be at risk of local extinction within their symbolic states by the end of the century (Table 2). For those species facing local extinction, some may find climatically suitable conditions in other states; however, many lack high-suitability areas near their current habitats, which may hinder natural dispersal and increase the need for assisted migration or targeted conservation efforts. While many state species may retain some suitable climatic habitat within their symbolic states, a large proportion are projected to experience significant declines in habitat suitability both locally and across the United States (Figure 5). In response to climate change, highly suitable climatic habitats for these species are expected to shift poleward and uphill, with this trend being more pronounced for state flowers than for state insects (Figure 4). These shifts threaten the symbolic identity of state species, disrupt local ecological interactions, and may lead to the loss of important ecosystem services.

### Beyond ecology and economics: cultural loss by climate-driven extinctions

Our findings (Figures 1-2) clearly show that numerous state flowers and insects are expected to undergo a significant reduction in areas with favorable climate within their states, with some experiencing such severe declines that it may result in dire consequences. Although a number of these species are not indigenous to their states and may not hold much economic or ecological importance, their potential disappearance poses a threat to states’ cultural heritage. It is challenging to quantify, let alone summarize, the cultural impacts of climate change because of the unique and diverse reasons behind the selection of each state species. However, a few anecdotes can help to illustrate some of the significant cultural losses at stake.

Minnesota’s state flower, the showy lady’s slipper (*C. reginae*), is projected to go from a very high median HSV of about 0.87 currently to a median value close to zero (∼ 0.04 by the 2080s, SSP5-8.5; Figure 1B), and will thus be at high risk of local extinction. This flower has been featured on the state flag since 1893 and was commemorated with the Minnesota Wildflower Route, which designated 130 km of State Highway 11 in its honor, where it is estimated that *>* 800, 000 of these flowers can be found today [35]. Our analysis suggests that this area will be climatically unsuitable for this species by the 2080s. Washington’s state flower, the rhododendron (*Rhododendron macrophyllum*), also faces a serious risk of local extinction. It was chosen through an 1893 vote, which was only open to women, 17 years before women were legally allowed to vote in elections [1].

Similarly, Oklahoma’s state flower, the mistletoe (*Phoradendron leucarpum*), was selected 14 years *before* statehood and is the oldest of the state’s symbols. Legend has it that the ability of the plant to withstand harsh winters symbolized the mettle of Oklahoma’s early pioneers [1]. Ironically, it may be warming winters that lead to the plant’s downfall and the loss of the state’s cultural heritage.

Montana’s bitterroot (*Lewisia rediviva*) is also endangered by climate change. The genus name honors Captain Meriwether Lewis of the Lewis and Clark Expedition. This historical journey holds significant cultural importance for the state (https://bit.ly/3yiu2co). The Bitterroot Mountains, Bitterroot Valley, and Bitterroot River owe their names to this state flower. Mary Long Anderson, who organized the referendum that selected the bitterroot in 1894, long before the state allowed women to vote, went on to become the state’s first congresswoman in 1916 [1].

These examples highlight the diverse and difficult-to-quantify cultural losses that may result from local extinctions caused by climate change. Similar to losing an important work of art, local loss of these species can have impacts that extend far beyond their ecological or economic effects [36].

### Potential ecological chain reactions triggered by habitat loss/ shifts of state species

Designating these flowers and insects as state species and implementing conservation policies has elevated their ecological importance within their symbolic states by supporting key functions such as pollination, host-plant interactions, and trophic relationships. However, many of these species are projected to experience declining habitat suitability—and in some cases, local extinction—within their designated states as climate change drives their distributions poleward and upward in elevation. For the 13 state species at risk of local extinction, even though suitable climatic habitats may emerge elsewhere, the geographic disconnect between current and future suitable areas may prevent natural migration without human intervention. These climate-driven range shifts could trigger cascading ecological consequences within local ecosystems, disrupting interactions and processes far beyond the loss of individual species [37, 38].

Among these, some species like the showy lady’s slipper (*C. reginae*), the mountain Laurel (*Kalmia latifolia*) and the large-flowered Trillium (*Trillium grandiflorum*) depend on mycorrhizal fungi for germination, nutrient uptake, and long-term survival [39, 40]. Climate-driven habitat loss may disrupt these specialized plant–fungus symbioses, potentially leading to the decline or local extinction of associated mycorrhizal fungi. Such disruptions can reduce fungal diversity and impair critical soil ecological functions, including nutrient cycling and plant community stability.

Some stress-tolerant plant species, such as Montana’s bitterroot *L. rediviva* and Tennessee coneflower *Echinacea tennesseensis*, act as ecological keystones in environments characterized by harsh and nutrient-poor conditions. The loss or decline of their climatic habitats can hinder vegetation recovery and compromise the overall resilience of these ecosystems. Their disappearance may also trigger cascading effects, including the collapse of local pollinator populations and disrupted reproduction in co-occurring plant species [37]. Furthermore, these species can be treated as bioindicators, the loss of which would diminish our capacity to monitor ecosystem health.

Species such as the two-spotted lady beetle (*Adalia bipunctata*) and the mourning cloak butterfly (*Nymphalis antiopa*), which function as native biocontrol agents and early-season pollinators, respectively, are susceptible to ecological displacement by invasive species or phenological mismatches. Their absence from symbolic states can destabilize local food web dynamics. The decline of *A. bipunctata*, for example, reduces natural aphid suppression and increases reliance on chemical pesticides, with downstream effects on non-target species and pollinators. The loss of *N. antiopa* may create a pollination gap for early-blooming plants, disrupting seasonal resource availability and reproductive cycles.

In summary, climate-driven range shifts among state species threaten not only the cultural heritage they represent but also the integrity and resilience of the ecosystems they support. Coordinated conservation efforts will be essential to prevent cascading ecological consequences and to safeguard both biodiversity and cultural heritage for future generations.

### Forward-looking conservation and management for preserving cultural heritage

As some state species may no longer find suitable climatic habitat within their symbolic states with climate change, traditional conservation approaches based solely on historical distributions may prove insufficient. Conservation planning should incorporate forward-looking strategies that account for projected habitat shifts [11].

#### Integrating climate adaptation, landscape connectivity, and community action

For endangered species such as the showy lady’s slipper (*C. reginae*), Tennessee coneflower (*E. tennesseensis*), and Karner blue butterfly (*Plebejus samuelis*), climate change is projected to reduce climatic habitat suitability to varying degrees. However, these species also face multiple stressors beyond climate, making their conservation especially complex. Identifying both remaining and newly emerging suitable climatic habitats under future climate scenarios is essential for guiding effective interventions.

In the case of the showy lady’s slipper, habitat loss from land conversion and environmental degradation currently poses a more immediate threat than climate itself. Conservation measures such as restoring alkaline wetlands and designating protected areas may help support this species [41]. However, such efforts may be insufficient in isolation, as future projections indicate a drastic decline in habitat suitability, potentially approaching zero.

Similarly, the Karner blue butterfly has experienced population declines due to both habitat loss and extreme weather events, such as the 2012 drought and heatwave [42]. In New Hampshire, its median habitat suitability is projected to fall from 0.85 to 0.25 under future climate conditions. Sustaining its populations will require targeted habitat management and safeguarding its sole larval host plant, wild blue lupine (*Lupinus perennis*).

To support the long-term persistence of these endangered species, conservation strategies must go beyond traditional habitat protection and incorporate climate-adaptive planning, landscape connectivity, and active stakeholder engagement.

#### From place-based strategy to dynamic, range-aware approaches

Place-based strategies have long been foundational in natural resource management and biodiversity conservation [11]. However, as climate change reshapes species distributions and alters ecosystem functions, static, localized conservation policies may no longer suffice. Species once tightly linked to particular geographies may face local extinction or shift outside their historical ranges, rendering traditional state or region-based protection strategies inadequate.

To address this, conservation planning must evolve toward dynamic, range-aware approaches that account for both current and projected future distributions. Regional management frameworks should be expanded and generalized to accommodate species’ anticipated movements, particularly along latitudinal and elevational gradients, where poleward and uphill shifts are increasingly common. This includes enhancing habitat connectivity to facilitate natural dispersal, especially across ecological corridors that align with these directional trends.

For state species facing significant climate-driven range contraction, suitable habitats may persist only in fragmented or limited areas within their symbolic states. Identifying these refugia and designating them as climate-resilient conservation zones is critical for maintaining ecological and cultural continuity. In cases where in-state persistence becomes unfeasible, coordinated inter-state conservation planning may be required to safeguard species in newly suitable habitats beyond their current political boundaries.

#### Replace with climate-resilient species

Alternative representative species may need to be considered to replace those originally selected for their popularity but now facing a high risk of local extinction due to climate change. Our findings suggest that certain taxonomic groups, such as shrubs and butterflies, may be more vulnerable to climate impacts than others (Figures 1C and 2C), indicating that these groups may be less ideal choices for long-term symbolic representation.

Some local state movements are dedicated to promoting the cultural identity of native species to align with a state’s priorities of maintaining local ecosystem stability. For example, movements in Alabama and Georgia are pushing to replace their current non-native state flowers with native alternatives [43, 44]. Although not all non-native species are inherently harmful—and some offer ecological or economic benefits [45, 46]—many pose significant risks to biodiversity, ecosystem function, and human health [47]. Moreover, managing non-native species is often complex due to their unpredictable ecological interactions.

Climate change adds further complexity to the native vs. non-native debate. While native species are more likely to face range contractions and local extinction, some non-native species may prove more resilient [48, 49]. Our results echo this concern, showing that non-native state species may be less vulnerable to climate impacts than their native counterparts. This raises a challenging dilemma: Should future state symbols prioritize climate resilience over ecological implications? As climate change accelerates, the debate over native versus non-native representation is likely to gain increasing attention in both conservation and cultural spheres.

### Limitations and future directions

Using species distribution models to assess the environmental niches of species based on their current distributions, combined with predicted future climate data, is the most convenient and effective technical approach for evaluating and comparing the resilience of different species to future climate change. However, due to the current limitations in the development of SDMs, it is challenging to account for species’ adaptability to their environments, the complex interactions between species and ecosystems, and the specificity of different terrains and landscapes. To improve the robustness of future assessments, it is essential to complement SDMs with empirical studies, expert knowledge, and Indigenous ecological perspectives. Such interdisciplinary integration can enhance our ability to identify viable alternative species or conservation strategies that are both ecologically appropriate and culturally meaningful.

Our study includes projections under two emissions scenarios but primarily focuses on the extreme high-emission scenario (SSP5-8.5) in the main analysis. As expected, the low-emissions scenario (SSP1-2.6) results in smaller quantitative impacts on habitat suitability for most species compared to SSP5-8.5. However, the qualitative patterns—such as the direction of range shifts and relative vulnerability—are broadly consistent between the two scenarios for both flowers and insects (Figure S3 and Figure 4A). Emissions scenarios are used in climate change research because future greenhouse gas trajectories are highly dependent on uncertain human behaviors. At present, global carbon emissions remain too high to meet international climate targets. Moreover, emerging evidence suggests that some forests [50, 51] and permafrost regions [52] may be transitioning from carbon sinks to carbon sources. These trends underscore a concerning outlook for future warming and its consequences for these cultural symbols. In this context, state species offer a compelling opportunity to engage the public in conversations about climate change and the urgent need for mitigation and adaptation strategies.

Overall, our findings contribute valuable insight into the climate vulnerability of culturally significant species. By highlighting their potential climatic habitat loss and range shifts, this work can raise public and policy-level awareness, foster support for adaptive conservation measures, and reinforce the societal importance of cultural identity, biodiversity, and ecosystem functionality.

## Supporting information

Supplementary Figuer S1-S3

## Acknowledgements

This work was conducted on the shared traditional territory of the Neutral, Anishnaabe and Haudenosaunee peoples. This land is part of the Dish with One Spoon Treaty between the Haudenosaunee and Anishnaabe peoples and symbolizes the agreement to share, protect our resources and not to engage in conflict. Acknowledging these nations reminds us of our important connection to the land and to our unfinished commitment to reconciliation. This work was supported by a grant to JAN from the Canadian Natural Science and Engineering Research Council (NSERC). The authors used ChatGPT (OpenAI, GPT-4), a large language model, to enhance writing clarity in some sections of the manuscript. All AI-assisted content was reviewed and verified by the authors.

## References

[1] Glynda Joy Nord. Official State Flowers and Trees. Trafford Publishing, Bloomington, Indiana, 2014.

[2] List of U.S. state and territory flowers. https://en.wikipedia.org/wiki/List_of_U.S._state_and_territory_flowers, 2023. (Accessed 12/04/2023).

[3] List of U.S. state insects. https://simple.wikipedia.org/wiki/List_of_U.S._state_insects, 2023. (Accessed 12/04/2023).

[4] List of state flowers and how they were chosen. https://www.jacksonandperkins.com/state-flowers-list-history/a/state-flowers-list-history/#:~:text=The%20diverse%20array%20of%20state,economic%20contributions%2C%20and%20regional%20identity., 2024. (Accessed on 07/23/2024).

[5] Rodrigo Cámara-Leret, N Raes, P Roehrdanz, Y De Fretes, CD Heatubun, L Roeble, A Schuiteman, PC Van Welzen, and L Hannah. Climate change threatens new guinea’s biocultural heritage. ScienceAdvances, 5(11):eaaz1455, 2019.

[6] Victoria Reyes-García, Rodrigo Cámara-Leret, Benjamin S Halpern, Casey O’hara, Delphine Renard, Noelia Zafra-Calvo, and Sandra Díaz. Biocultural vulnerability exposes threats of culturally important species. Proceedings of the National Academy of Sciences, 120(2):e2217303120, 2023.

[7] Magdalena Pasikowska-Schnass. The impact of climate change on cultural heritage. Technical report, European Parliamentary Research Service, 2024.

[8] I-Ching Chen, Jane K Hill, Ralf Ohlemüller, David B Roy, and Chris D Thomas. Rapid range shifts of species associated with high levels of climate warming. Science, 333(6045):1024–1026, 2011.

[9] John J Wiens. Climate-related local extinctions are already widespread among plant and animal species. PLoS biology, 14(12):e2001104, 2016.

[10] Gretta T Pecl, Rachel Kelly, Chloe Lucas, Ingrid van Putten, Renuka Badhe, Curtis Champion, I-Ching Chen, Omar Defeo, Juan Diego Gaitan-Espitia, Birgitta Evengård, et al. Climate-driven ‘species-on-the-move’provide tangible anchors to engage the public on climate change. People and Nature, 5(5):1384–1402, 2023.

[11] Jonathan W Moore and Daniel E Schindler. Getting ahead of climate change for ecological adaptation and resilience. Science, 376(6600):1421–1426, 2022.

[12] Gian-Reto Walther. Community and ecosystem responses to recent climate change. Philosophical Transactions of the Royal Society B: Biological Sciences, 365(1549):2019–2024, 2010.

[13] Scott Chamberlain, Vijay Barve, Dan Mcglinn, Damiano Oldoni, Peter Desmet, Laurens Geffert, and Karthik Ram. rgbif: Interface to the Global Biodiversity Information Facility API, 2023. R package version 3.7.8.

[14] Xuezhen Ge, Cortland K Griswold, and Jonathan A Newman. Robust species distribution predictions of predator and prey responses to climate change. Journal of Biogeography, 2024.

[15] Alexander Zizka, Daniele Silvestro, Tobias Andermann, Josué Azevedo Camila Duarte Ritter, Daniel Edler, Harith Farooq, Andrei Herdean, María Ariza, Ruud Scharn, et al. Coordinatecleaner: Standardized cleaning of occurrence records from biological collection databases. Methods in Ecology and Evolution, 10(5):744–751, 2019.

[16] Roeland Kindt and Richard Coe. Tree diversity analysis. A manual and software for common statistical methods for ecological and biodiversity studies, 2005. ISBN 92-9059-179-X.

[17] Morgane Barbet-Massin, Frédéric Jiguet, Cécile Hélène Albert, and Wilfried Thuiller. Selecting pseudoabsences for species distribution models: how, where and how many? Methods in Ecology and Evolution, 3(2):327–338, 2012.

[18] Wilfried Thuiller, Damien Georges, Maya Gueguen, Robin Engler, Frank Breiner, Bruno Lafourcade, and Remi Patin. biomod2: Ensemble Platform for Species Distribution Modeling, 2023. R package version 4.2-3.

[19] Roozbeh Valavi, Jane Elith, José J Lahoz-Monfort, and Gurutzeta Guillera-Arroita. blockCV: An R package for generating spatially or environmentally separated folds for k-fold cross-validation of species distribution models. Methods in Ecology and Evolution, 10(2):225–232, 2019.

[20] Dirk Nikolaus Karger, Olaf Conrad, Jürgen Bohner, Tobias Kawohl, Holger Kreft, Rodrigo Wilber Soria-Auza, Niklaus E Zimmermann, H Peter Linder, and Michael Kessler. Climatologies at high resolution for the earth’s land surface areas. Scientific data, 4(1):1–20, 2017.

[21] Gregory Flato, Jochem Marotzke, Babatunde Abiodun, Pascale Braconnot, S Chan Chou, William Collins, Peter Cox, Fatima Driouech, Seita Emori, Veronika Eyring, et al. Evaluation of climate models. In Climate change 2013: the Physical Science Basis. Contribution of Working Group I to the Fifth Assessment Report of the Intergovernmental Panel on Climate Change, pages 741–866. Cambridge University Press, 2014.

[22] Suchada Kamworapan and Chinnawat Surussavadee. Evaluation of CMIP5 global climate models for simulating climatological temperature and precipitation for Southeast Asia. Advances in Meteorology, 2019:1–18, 2019.

[23] Peter McCullagh and John A Nelder. Generalized linear models. Chapman and Hall, 2 edition, 1989.

[24] Trevor Hastie and Robert Tibshirani. Generalized additive models. Statistical Science, 1(3):297–310, 1986.

[25] Greg Ridgeway. The state of boosting. Computing Science and Statistics, pages 172–181, 1999.

[26] Jerome H Friedman. Multivariate adaptive regression splines. The Annals of Statistics, 19(1):1–67, 1991.

[27] Leo Breiman. Classification and regression trees. Routledge, New York, 1 edition, 1984.

[28] Trevor Hastie, Robert Tibshirani, and Andreas Buja. Flexible discriminant analysis by optimal scoring. Journal of the American Statistical Association, 89(428):1255–1270, 1994.

[29] Steven J Phillips, Robert P Anderson, and Robert E Schapire. Maximum entropy modeling of species geographic distributions. Ecological Modelling, 190(3-4):231–259, 2006.

[30] Wilfried Thuiller, Sandra Lavorel, Miguel B Araújo, Martin T Sykes, and I Colin Prentice. Climate change threats to plant diversity in europe. Proceedings of the National Academy of Sciences, 102(23):8245–8250, 2005.

[31] Miguel B Araújo, Wilfried Thuiller, and Richard G Pearson. Climate warming and the decline of amphibians and reptiles in europe. Journal of biogeography, 33(10):1712–1728, 2006.

[32] Canran Liu, Matt White, and Graeme Newell. Selecting thresholds for the prediction of species occurrence with presence-only data. Journal of biogeography, 40(4):778–789, 2013.

[33] Cândida G Vale, Pedro Tarroso, and José C Brito. Predicting species distribution at range margins: testing the effects of study area extent, resolution and threshold selection in the sahara–sahel transition zone. Diversity and Distributions, 20(1):20–33, 2014.

[34] Ya Zou, Xuezhen Ge, Shixiang Zong, and Jonathan A Newman. Climate change may make pine wilt disease more prevalent. Journal of Applied Ecology, 61(12):3028–3039, 2024.

[35] Scenic drives northern mn. https://visitwarroad.com/scenic-drives/#:~:text=The%20Wildflower%20Route%20on%20Highway,along%20the%2075%2Dmile%20stretch., 2024. (Accessed on 07/24/2024).

[36] Jonathan A. Newman, Gary Varner, and Stefan Linquist. Defending Biodiversity: Environmental Science and Ethics. Cambridge University Press, 2017.

[37] Jane Memmott, Paul G Craze, Nickolas M Waser, and Mary V Price. Global warming and the disruption of plant–pollinator interactions. Ecology letters, 10(8):710–717, 2007.

[38] Jason M Tylianakis, Raphael K Didham, Jordi Bascompte, and David A Wardle. Global change and species interactions in terrestrial ecosystems. Ecology letters, 11(12):1351–1363, 2008.

[39] DJ Read. The structure and function of the ericoid mycorrhizal root. Annals of botany, 77(4):365–374, 1996.

[40] Marcel GA Van Der Heijden, Richard D Bardgett, and Nico M Van Straalen. The unseen majority: soil microbes as drivers of plant diversity and productivity in terrestrial ecosystems. Ecology letters, 11(3):296–310, 2008.

[41] Cypripedium reginae. https://en.wikipedia.org/wiki/Cypripedium_reginae, 2024. (Accessed on 07/30/2024).

[42] Karner blue. https://en.wikipedia.org/wiki/Karner_blue, 2024. (Accessed on 07/30/2024).

[43] Time to change the state flower from cherokee rose to a native plant. https://georgiarecorder.com/2024/02/26/time-to-change-the-state-flower-from-cherokee-rose-to-a-native-plant/, 2024. (Accessed on 07/30/2024).

[44] Now is your chance to change the state flower of georgia to a native species! https://www.nurturenativenature.com/post/now-is-your-chance-to-change-the-state-flower-of-georgia-to-a-native-species, 2024. (Accessed on 07/30/2024).

[45] LS Wootton, SD Halsey, K Bevaart, A McGough, J Ondreicka, and P Patel. When invasive species have benefits as well as costs: managing carex kobomugi (asiatic sand sedge) in new jersey’s coastal dunes. Biological Invasions, 7:1017–1027, 2005.

[46] Melina Kourantidou, Phillip J Haubrock, Ross N Cuthbert, Thomas W Bodey, Bernd Lenzner, Rodolphe E Gozlan, Martin A Nun∼ez, Jean-Michel Salles, Christophe Diagne, and Franck Courchamp. Invasive alien species as simultaneous benefits and burdens: trends, stakeholder perceptions and management. Biological Invasions, 24(7):1905–1926, 2022.

[47] Prabhat Kumar Rai and JS Singh. Invasive alien plant species: Their impact on environment, ecosystem services and human health. Ecological indicators, 111:106020, 2020.

[48] Heather A Hager, Geraldine D Ryan, Hajnal M Kovacs, and Jonathan A Newman. Effects of elevated co<sub>2 </sub>on photosynthetic traits of native and invasive c3 and c4 grasses. BMC ecology, 16:1–13, 2016.

[49] Bethany A Bradley, Evelyn M Beaury, Belinda Gallardo, Inés Ibán∼ez, Catherine Jarnevich, Toni Lyn Morelli, Helen R Sofaer, Cascade JB Sorte, and Montserrat Vila. Observed and potential range shifts of native and nonnative species with climate change. Annual Review of Ecology, Evolution, and Systematics, 55, 2024.

[50] Matt McGrath. World way off target in tackling climate change - UN. https://www.bbc.com/news/articles/ce8yyle2eq2o, 2024. (Accessed on 11/14/2024).

[51] Ben Turner. Global carbon emissions reach new record high in 2024, with no end in sight, scientists say. https://www.livescience.com/planet-earth/climate-change/global-carbon-emissions-reach-new-record-high-in-2024-with-no-end-in-sight-scientists-say, 2024. (Accessed on 11/14/2024).

[52] Claire C. Treat, Anna-Maria Virkkala, Eleanor Burke, Lori Bruhwiler, Abhishek Chatterjee, Joshua B. Fisher, Josh Hashemi, Frans-Jan W. Parmentier, Brendan M. Rogers, Sebastian Westermann, Jennifer D. Watts, Elena Blanc-Betes, Matthias Fuchs, Stefan Kruse, Avni Malhotra, Kimberley Miner, Jens Strauss, Amanda Armstrong, Howard E. Epstein, Bradley Gay, Mathias Goeckede, Aram Kalhori, Dan Kou, Charles E. Miller, Susan M. Natali, Youmi Oh, Sarah Shakil, Oliver Sonnentag, Ruth K. Varner, Scott Zolkos, Edward A.G. Schuur, and Gustaf Hugelius. Permafrost carbon: Progress on understanding stocks and fluxes across northern terrestrial ecosystems. Journal of Geophysical Research: Biogeosciences, 129(3):e2023JG007638, 2024.

