## Supplement 1 for "Wilting Wildflowers and Bummed-Out Bees: Climate Change Threatens U.S. State Symbols"

**Supplement 1. Detailed documentation of the state flower species examined in this study.**

| State | Category | Common name (Scientific name) | Notes |
|---|---|---|---|
| Alabama (AL) | state flower | Common camellia or Japanese camellia (*Camellia japonica*) | **Evidence of designating as the official state flower:**<br>● From The Code of Alabama, Title 1, Chapter 2, Section 1-2-11. "The camellia, *Camellia japonica* L., is hereby designated and named as the official state flower of Alabama." [AL-1] |
| | state wildflower | Oak-leaf hydrangea (*Hydrangea quercifolia*) | **Evidence of designating as the official state wildflower:**<br>● "Alabama designated the oak-leaf hydrangea (*Hydrangea quercifolia* Bartr.) as the official state wildflower in 1999."[AL-2] |
| Alaska (AK) | state flower | Alpine forget-me-not (*Myosotis asiatica*) | **Evidence of designating as the official state flower:**<br>● From The Alaska Statutes - 2004, Title 44, Chapter 09, Section 44-09-050. "The wild native forget-me-not is the state flower and floral emblem." [AK-1]<br>**Synonym(s):**<br>● *Myosotis asiatica*, *Myosotis alpestri*, *Myosotis alpestris ssp. asiatica*, *Myosotis sylvatica var. alpestris.*<br>● Raw GBIF records include the occurrence records searched by all the scientific names. |
| Arizona (AZ) | state flower | Saguaro cactus blossom (*Carnegiea gigantea*) | **Evidence of designating as the official state flower:**<br>● From the Arizona Revised Statutes, Title 41, Chapter 4.1, Article 5, Section 41-855. "The pure white waxy flower of the cereus giganteus (giant cactus) or Saguaro shall be the state flower." [AZ-1]<br>**Synonym(s):**<br>● *Carnegiea gigantea*, *Cereus giganteus*<br>● Raw GBIF records include the occurrence records searched by all the scientific names. |
| Arkansas (AR) | state flower | Apple blossom (*Malus domestica*) | **Evidence of designating as the official state flower:**<br>● From the Arkansas Code (Non annotated), Title 1, Chapter 2, Section 1-4-109. "The apple blossom is declared to be the state floral emblem of Arkansas." [AR-1]<br>● The legislation did not specify a specific species of the apple blossom and some people are referring it to as crabapple (*Malus coronaria*). However, we think it refers to *Malus domestica* because they mentioned apple industry, such as "On the day that the legislature was to vote, Mrs. Barton appeared at the Capitol in a red dress and a crisp red Arkansas apple for |

| | | | |
|---|---|---|---|
| | | | each legislator."[AR-1] and "Arkansas was once a major apple producing state and still has an Arkansas apple festival each year in the town of Lincoln (in Washington county)."[AR-2]<br>**Synonym(s):**<br>● *Malus domestica, Malus pumila*<br>● Raw GBIF records include the occurrence records searched by all the scientific names. |
| California (CA) | state flower | California poppy (*Eschscholzia californica*) | **Evidence of designating as the official state flower:**<br>● From the California Government Code, General Provisions, Title 1, Division 2, Section 421. "The golden poppy (Eschscholzia) is the official State Flower. April 6 of each year is hereby designated California Poppy Day." [CA-1]<br>● Though the act stopped short of naming a particular species, the California poppy (*Eschscholzia californica*) is the variety generally thought of as the official State flower. [CA-1] |
| Colorado (CO) | state flower | Colorado blue columbine (*Aquilegia coerulea*) | **Evidence of designating as the official state flower:**<br>● From the Colorado Revised Statutes, Title 1, Part 9, Sections 24-80-905 through 24-80-908. "The white and lavender columbine is hereby made and declared to be the state flower of the state of Colorado." [CO-1]<br>● The act names the white and lavender columbine as the state flower without reference to a scientific name. In later legislation declaring it the duty of the citizens of the state to protect the state flower, the white and lavender columbine is referred to as *Aquilegia caerulea*. [CO-1] |
| Connecticut (CT) | state flower | Mountain laurel (*Kalmia latifolia*) | **Evidence of designating as the official state flower:**<br>● From The General Statutes of Connecticut, Title 3, Chapter 3, Section 3-108. "The mountain laurel, *Kalmia latifolia*, shall be the state flower." [CT-1]<br>● The mountain laurel (*Kalmia latifolia*) was declared to be the state flower of the State of Connecticut by an act of the General Assembly approved on April 17, 1907. [CT-1]<br>**Same state flower:**<br>● *Kalmia latifolia* is the official state flower of Connecticut (CT) and Pennsylvania (PA). |
| | Children's state flower | Michaela Petit's four-o'clocks (*Mirabilis jalapa*) | **Evidence of designating as the official state flower:**<br>● "Michaela Petit's Four-O'Clocks, *Mirabilis jalapa*, shall be the children's state flower."[CT-2] |

| | | | |
|---|---|---|---|
| Delaware (DE) | State flower | Peach blossom (*Prunus persica*) | **Evidence of designating as the official state flower:**<br>● From The Delaware Code, Title 29, Chapter 3, Section 308. “The peach blossom, as originally adopted as the floral emblem of the State on May 9, 1895, shall be the official state flower. (29 Del. C. 1953, § 508; 50 Del. Laws, c. 289, § 1.)”[DE-1]<br>● “The peach blossom was adopted as Delaware's floral emblem by an act of the legislature on March 9, 1895. In 1953, it was named as the official state flower.”[DE-1] |
| Florida (FL) | state flower | Orange blossom (*Citrus sinensis*) | **Evidence of designating as the official state flower:**<br>● “The orange blossom (*Citrus sinensis*) was adopted by a Concurrent Resolution of the Florida state legislature on May 5, 1909.” [FL-1]<br>**Occurrence records:**<br>● From the GBIF, we obtained only 35 occurrence records in the U.S. Since the distribution of *Citrus sinensis* in the U.S. can also be represented by orange reproduction data, we downloaded U.S. citrus production statistics from https://quickstats.nass.usda.gov using the following term selections: Program: Census; Sector: Crops; Group: Fruit & Tree Nuts; Commodity: Oranges; Data Item: Oranges – Acres Bearing & Non-Bearing; Domain: Total; Geographic Level: County; Years: 2017 and 2012; Period Type: Annual. We obtained 214 coordinates and combined them with the GBIF records to represent the presence data for the species. |
| | state wildflower | Tickseed (*Coreopsis* spp.) | **Evidence of designating as the official state wildflower:**<br>● “In 1991 the flower of the genus *Coreopsis* was designated as Florida's official wildflower. The state legislature made this designation after the colorful flowers were used extensively in Florida's roadside plantings and highway beautification programs. The coreopsis is found in a variety of colors, ranging from golden to pink.” [FL-2] |
| Georgia (GA) | state flower | Cherokee rose (*Rosa laevigata*) | **Evidence of designating as the official state flower:**<br>● “The Cherokee rose was adopted by the Georgia General Assembly as the floral emblem of the State of Georgia at the request of the Federation of Women's Clubs. It was adopted by Joint Resolution No. 42 approved on August 18, 1916.” [GA-1]<br>● From the Georgia Code, Title 50, Chapter 3, Section 50-3-53. “The Cherokee rose is adopted as the floral emblem of the State of Georgia.” [GA-1] |

| | | | |
|---|---|---|---|
| | state wildflower | Azalea (*Rhododendron* spp.) | **Evidence of designating as the official state wildflower:**<br>● "Georgia designated azalea as the official state wildflower in 1979 (the state flower is the Cherokee rose). In 2013 the Act was amended to specify native azaleas (*Rhododendron* sp.), collectively, as Georgia's state wildflower symbol."[GA-2] |
| Idaho (ID) | state flower | Syringa, mock orange (*Philadelphus lewisii*) | **Evidence of designating as the official state flower:**<br>● From the Idaho Statutes, Title 67, Chapter 45, Section 67-4502. "The Syringa (*Philadelphus lewisii*) is hereby designated and declared to be the state flower of the state of Idaho." [ID-1] |
| Illinois (IL) | state flower | Common blue violet (*Viola sororia*) | **Evidence of designating as the official state flower:**<br>● "The law of 1908 designated that the "blue violet" be the state flower. There are actually eight species of blue-flowered violets in the state. The most common of them is the dooryard violet, *Viola sororia*. The dooryard violet is certainly one of the most recognizable native wildflowers in the state. It is also one of the most easily grown; it grows in anything from full sunlight to deep shade." [IL-1]<br>**Synonym(s):**<br>● *Viola sororia*, *Viola papilionacea*<br>● Raw GBIF records include the occurrence records searched by all the scientific names.<br>**Same state flower:**<br>● *Viola sororia* is the state flower of Illinois (IL), New Jersey (NJ), Rhode Island (RI), and Wisconsin (WI). |
| | state wildflower | Milkweed (*Asclepias* spp.) | **Evidence of designating as the official state wildflower:**<br>● The plant Asclepias spp, commonly known as "Milkweed", is designated the official State wildflower of the State of Illinois. [IL-2] |
| Indiana (IN) | state flower | Peony (*Paeonia* spp.) | **Evidence of designating as the official state flower:**<br>● From the Illinois Compiled Statutes, Government, Chapter 5, State Designations Act, Section 40. "The white oak tree is designated the native State tree of the State of Illinois; and the native violet is designated the native State flower of the State of Illinois." [IN-1]<br>● "The legislation did not specify a specific variety of violet but, according to the Illinois State Museum, the dooryard or common violet (*Viola sororia*) is the most common species in the state and was probably the intended "native violet" of Senator Jackson's Bill." [IN-1]<br>**Multiple species:** |

| | | | |
|---|---|---|---|
| | | | ● *Paeonia lactiflora* (common garden peony, native to Asian) and *Paeonia officinalis* (common peony, native to European) are most commonly bred into herbaceous cultivars, so we modelled the potential distributions for two species in this state. |
| | state wildflower | Milkweed, butterflyweed (*Asclepias tuberosa*) | **Evidence of designating as the official state wildflower:**<br>● "Indiana has an Official Firearm, and yes, an Official State Pie. It also has a State Flower, of course—the peony. And now it may get a new State Wildflower, that vibrant orange pollinator-and-monarch magnet, Butterfly Milkweed, aka Butterflyweed (*Asclepias tuberosa*)." [IN-2]<br>● Although *Asclepias tuberosa* proposed to be the state wildflower of Indiana, it looks like it wasn't approved by legislation [IN-3] [IN-4]. However, we think it is worth to include it in our study for future references. |
| Kansas (KS) | state flower | Wild native sunflower (*Helianthus annuus*) | **Evidence of designating as the official state flower:**<br>● From the Kansas Statutes, Chapter 73, Article 18, Section 73-1801. "Be it enacted by the Legislature of the State of Kansas: That the helianthus or wild native sunflower is hereby made, designated and declared to be the state flower and floral emblem of the state of Kansas." [KS-1] |
| Kentucky (KY) | state flower | Goldenrod (*Solidago gigantea*) | **Evidence of designating as the official state flower:**<br>● From the Kentucky Revised Statutes, Title 1, Chapter 2, Section 2.09. "The goldenrod is the official state flower of Kentucky." [KY-1]<br>● "The website of Kentucky Department for Libraries and Archives names *Solidago gigantea* as the state flower." [KY-1]<br>**Same state flower:**<br>● *Solidago gigantea* is the state flower of Kentucky (KY) and Nebraska (NE). |
| Louisiana (LA) | state flower | Southern magnolia (*Magnolia grandiflora*) | **Evidence of designating as the official state flower:**<br>● From the Louisiana Revised Statutes, Title 49, Part 8, Section RS 49:154. "The magnolia shall be the state flower of the State of Louisiana. Acts 1990, No. 511, §1." [LA-1]<br>● "Louisiana by legislative action, approved July 12, 1900, designated the magnolia [*Magnolia foetida* or *Magnolia grandiflora*] as the State flower. It was chosen probably because there is such an abundant growth of this tree throughout the State." [LA-1]<br>**Synonym(s):**<br>● *Magnolia grandiflora*, *Magnolia foetida*<br>● Raw GBIF records include the occurrence records searched by all the scientific names |

| | | | |
|---|---|---|---|
| | | | **Same state flower:**<br>● *Magnolia grandiflora* is the state flower of Louisiana (LA) and Mississippi (MS). |
| | state wildflower | Louisiana iris (*Iris giganticaerulea*) | **Evidence of designating as the official state wildflower:**<br>● "The Louisiana iris (*Iris giganticaerulea*) was designated the official state wildflower in 1990."[LA-2] |
| Iowa (IA) | state flower | Wild rose (*Rosa arkansana*) | **Evidence of designating as the official state flower:**<br>● "The wild rose was adopted by resolution of the Iowa General Assembly and is not included as statutory law in the Iowa Code." [IA-1]<br>● "A specific variety of wild rose was not named in the legislation though *Rosa pratincola* (Synonym: *Rosa arkansana*) is thought to represent the variety intended by the legislation. The wild rose, adopted by the concurrent resolution, is often referred to as the wild prairie rose today." [IA-1] |
| Maine (ME) | state flower | White pine cone and tassel (*Pinus strobus*) | **Evidence of designating as the official state flower:**<br>● From the Maine Revised Statutes, Title 1, Chapter 9, Subchapter 1, Section 211. "The floral emblem for the State, in the national garland of flowers, shall be the pine cone and tassel. " [ME-1]<br>● Though not named in the act of the legislature, the pine cone and tassel intended by the legislation is that of the eastern white pine (*Pinus strobus*). [ME-1] |
| Maryland (MD) | state flower | Black-eyed Susan (*Rudbeckia hirta*) | **Evidence of designating as the official state flower:**<br>● From the Maryland Statutes, Title 13, Section 13-305. "The black-eyed susan (*Rudbeckia hirta*) is the State flower." [MD-1] |
| Massachusetts (MA) | state flower | Mayflower (*Epigaea repens*) | **Evidence of designating as the official state flower:**<br>● From the General Laws of Massachusetts, Part 1, Title 1, Chapter 2, Section 7. "The mayflower (*Epigaea repens*) shall be the flower or floral emblem of the commonwealth. It was amended to protect the endangered mayflower." [MA-1] |
| Michigan (MI) | state flower | Apple blossom (*Malus coronaria* and *Malus domestica*) | **Evidence of designating as the official state flower:**<br>● "WHEREAS, Our blossoming apple trees add much to the beauty of our landscape, and Michigan apples have gained a worldwide reputation; and WHEREAS, At least one of the most fragrant and beautiful flowered species of apple, the pyrus coronaria, is native to our state; therefore Resolved by the Senate and House of Representatives of the State of Michigan, That the apple blossom be and the same hereby is designated and adopted as the state flower of the state of Michigan." [MI-1] |

| | | | |
|---|---|---|---|
| | | | ● "Citing the blossom of the native Michigan *Pyrus coronaria* (sweet crabapple) as particularly beautiful and fragrant, the legislation does not specify this species as the state flower but refers to the generic apple blossom as the state flower of Michigan." [MI-1]<br>● In the legislation, the preamble refers to both the native crab apple (Pyrus/Malus coronaria) and says "... and Michigan apples have gained a worldwide reputation", which would have to refer to the apple industry, i.e., *Malus domestica*. Therefore, we include both of the *Malus coronaria* and *Malus domestica* in our study.<br>**Synonym(s):**<br>● *Malus domestica, Malus pumila*<br>● Raw GBIF records include the occurrence records searched by all the scientific names. |
| | state wildflower | Dwarf lake iris *(Iris lacustris)* | **Evidence of designating as the official state wildflower:**<br>● "In 1998, the DWARF LAKE IRIS (*Iris lacustris*) was designated as the state wildflower. Native to the state, the endangered flower grows along the northern shorelines of Lakes Michigan and Huron." [MI-2] |
| Minnesota (MN) | state flower | Pink and white lady's slipper (*Cypripedium reginae*) | **Evidence of designating as the official state flower:**<br>● From the Minnesota Statutes, Chapter 1, Section 1.142 and Chapter 18H, Section 18H.18. "The pink and white lady slipper, *Cypripedium reginae*, is the official flower of the state of Minnesota." [MN-1] |
| Mississippi (MS) | state flower | Southern magnolia (*Magnolia grandiflora*) | **Evidence of designating as the official state flower:**<br>● From the Mississippi Code, Title 3, Chapter 3, Section 3-3-13. The flower or bloom of the magnolia or evergreen magnolia (*Magnolia grandiflora* L.) is hereby designated as the state flower of Mississippi. [MS-1]<br>**Same state flower:**<br>● *Magnolia grandiflora* is the state flower of Louisiana (LA) and Mississippi (MS). |
| | state wildflower | Tickseed (*Coreopsis* spp.) | **Evidence of designating as the official state wildflower:**<br>● "Mississippi designated coreopsis as the official state wildflower in 1991."[MS-2] |
| Missouri (MO) | state flower | Downy hawthorn *(Crataegus mollis*) | **Evidence of designating as the official state flower:**<br>● From the Missouri Revised Statutes, Title 2, Chapter 10, Section 10.030. "The hawthorn, the blossom of the tree commonly called the "red haw" or "wild haw" and scientifically designated as crataegus, is declared to be the floral emblem of Missouri, and the state department of agriculture shall |

| | | | |
|---|---|---|---|
| | | | recognize it as such and encourage its cultivation on account of the beauty of its flower, fruit and foliage.” [MO-1]<br>● “Though a specific variety of hawthorn is not named in the legislation, the Missouri Department of Conservation asserts that the downy hawthorn (*Crataegus mollis*) is the species deserving of the recognition.” [MO-1] |
| Montana (MT) | state flower | Bitterroot (*Lewisia rediviva*) | **Evidence of designating as the official state flower:**<br>● From the Montana Code, Title 1, Chapter 1, Part 5, Section 1-1-503. “The flower known as *Lewisia rediviva* (bitterroot) shall be the floral emblem of the state of Montana.” [MT-1] |
| Nebraska (NE) | state flower | Goldenrod (*Solidago gigantea*) | **Evidence of designating as the official state flower:**<br>● We, the Legislature of Nebraska hereby declare the flower commonly known as the "Golden Rod" (*Solidago serotina*) to be the floral emblem of the state. [NE-1]<br>● The flower, spelled "Golden Rod" in the legislation and referred to as "*Solidago serotina*" is commonly called giant goldenrod (*Solidago gigantea*) today. [NE-1]<br>**Same state flower:**<br>*Solidago gigantea* is the state flower of Kentucky (KY) and Nebraska (NE). |
| Nevada (NV) | state flower | Sagebrush (*Artemisia tridentata*) | **Evidence of designating as the official state flower:**<br>● From the Nevada Revised Statutes, Title 19, Chapter 235, Section 235.050. “The shrub known as Sagebrush (*Artemisia tridentata* or *trifida*) is hereby designated as the official state flower of the State of Nevada.” [NV-1]<br>● *Artemisia trifida* only has one occurrence record in GBIF, so we only chose *Artemisia tridentata* as the state flower in our study. |
| New Hampshire (NH) | state flower | Purple lilac (*Syringa vulgaris*) | **Evidence of designating as the official state flower:**<br>● From from the New Hampshire Revised Statutes, Title 1, Chapter 3, Section 3:5. “The purple lilac, *Syringa vulgaris*, is the state flower of New Hampshire.” [NH-1] |
| | state wildflower | Pink lady’s slipper (*Cypripedium acaule*) | **Evidence of designating as the official state wildflower:**<br>● “The pink lady’s slipper, *Cypripedium acaule*, is hereby designated as the official state wildflower of New Hampshire.” [NH-2]<br>● “In 1991, the Pink Lady’s Slipper became the state’s wildflower. The plant is native to New Hampshire and grows in the moist wooded areas of the state.” [NH-2] |

| | | | |
|---|---|---|---|
| New Jersey (NJ) | state flower | Common blue violet (Viola sororia) | **Evidence of designating as the official state flower:**<br>● From the New Jersey Permanent Statutes, Title 52, Section 52:9A-2. “The violet (common meadow, *V. sororia*) is designated the New Jersey State Flower.” [NJ-1]<br>**Synonym(s):**<br>● *Viola sororia*, *Viola papilionacea*<br>● Raw GBIF records include the occurrence records searched by all the scientific names.<br>**Same state flower:**<br>● *Viola sororia* is the state flower of Illinois (IL), New Jersey (NJ), Rhode Island (RI), and Wisconsin (WI). |
| New Mexico (NM) | state flower | Yucca (*Yucas* spp.) | **Evidence of designating as the official state flower:**<br>● From the New Mexico Statutes, Article 3, Section 12-4-4 A. “The yucca flower is adopted as the official flower of New Mexico.” [NM-1]<br>● “The legislation does not specify a particular species of yucca flower or even indicate that "all" species are intended to represent New Mexico.” “We have not been able to locate any credible evidence that a particular species of yucca was intended as the state flower and until we do, we have to assume that any species may be considered to be the "official Flower of the State of New Mexico." [NM-1]<br>**Multiple species:**<br>● There are 27 Yucas spp. in the United States. 11 of them are found in New Mexico, there are Yucca baccata, Yucca campestris, Yucca elata, Yucca faxoniana, Yucca glauca, Yucca harrimaniae, Yucca madrensis, Yucca schottii, Yucca torreyi, Yucca baileyi [NM-2]. So we modelled the potential distribution for the 11 species. |
| North Carolina (NC) | state flower | Flowering dogwood (*Cornus florida*) | **Evidence of designating as the official state flower:**<br>● From the North Carolina General Statutes, Chapter 145, Section 145-1. “The dogwood is hereby adopted as the official flower of the State of North Carolina. (1941, c. 289.)” [NC-1]<br>● “Though not specified in the legislation, *Cornus florida*, commonly referred to as the flowering dogwood, is accepted as the species intended as the official flower of North Carolina.” [NC-1] |
| | state wildflower | Carolina lily (*Lilium michauxii*) | **Evidence of designating as the official state wildflower:**<br>● “North Carolina designated the Carolina lily (*Lilium michauxii*) as the official State wildflower in 2003. This spectacular wildflower grows throughout |

| | | | |
|---|---|---|---|
| | | | North Carolina, from the forests and hills of Cherokee County to the coastal swamplands of Hyde and Pamlico Counties." [NC-2] |
| North Dakota (ND) | state flower | Wild prairie rose (*Rosa arkansana* and *Rosa blanda*) | **Evidence of designating as the official state flower:**<br>● From the North Dakota Century Code, Title 54, Chapter 54-02, Section 54-02-03. "The floral emblem of the state of North Dakota shall be the wild prairie rose, *Rosa blanda* or *arkansana*." [ND-1]<br>**Multiple species:**<br>● We model the potential distributions for the two species in this state. |
| Ohio (OH) | state wildflower | Large white trillium (*Trillium grandiflorum*) | **Evidence of designating as the official state wildflower:**<br>● "In 1986, the Ohio General Assembly made the white trillium Ohio's official wildflower…… The General Assembly selected this flower because it exists in all of Ohio's 88 counties." [OH-1] |
| Oregon (OR) | state flower | Oregon grape (*Mahonia aquifolium*) | **Evidence of designating as the official state flower:**<br>● "According to Senate Concurrent Resolution Number Four of the Oregon Legislative Assembly, the State of Oregon on January 30-31, 1899, adopted as her State flower, the Oregon grape (*Berberis aquifolium*)." [OR-1]<br>**Synonym(s):**<br>● *Mahonia aquifolium, Berberis aquifolium*<br>● Raw GBIF records include the occurrence records searched by all the scientific names. |
| Oklahoma (OK) | state floral emblem | Mistletoe (*Phoradendron leucarpum*) | **Evidence of designating as the official floral emblem:**<br>● "A parasitic plant used in Christmas decorations, mistletoe (*Phoradendron serotinum*) is Oklahoma's official floral emblem. The territorial legislature so designated the plant on February 11, 1893, in House Bill 49. Introduced by Rep. J. A. Wimberly. The bill was the first to establish an official floral emblem by any legislature." [OK-1] |
| | state wildflower | Indian blanket (*Gaillardia pulchella*) | **Evidence of designating as the official state wildflower:**<br>● "The "Indian Blanket" was approved as Oklahoma's official state wildflower in 1986." [OK-2] |
| Pennsylvania (PA) | state flower | Mountain laurel (*Kalmia latifolia*) | **Evidence of designating as the official state flower:**<br>● From Thomas E. Martin, Jr., Esq.'s Unconsolidated Pennsylvania Statutes, Title 71. "The mountain laurel (*Kalmia Latifolia*) is hereby adopted as the State flower of Pennsylvania." [PA-1]<br>**Same state flower:** |

| | | | |
|---|---|---|---|
| | | | ● *Kalmia latifolia* is the official state flower of Connecticut (CT) and Pennsylvania (PA). |
| | beautification and conservation plant | Penngift crown vetch (*Coronilla varia*) | **Evidence of designating as the official state beautification and conservation plant:**<br>● "Declaring and adopting Penngift Crownvetch (*Coronilla varia* L. Penngift) as the State Beautification and Conservation Plant of Pennsylvania." [PA-2] |
| Rhode Island (RI) | state flower | Common blue violet (Viola sororia) | **Evidence of designating as the official state flower:**<br>● From the State of Rhode Island General Laws, Title 42, Chapter 42-4, Section 42-4-9. "The flower commonly known as the "violet" (*Viola sororia*) is hereby designated as the state flower." [RI-1]<br>**Synonym(s):**<br>● *Viola sororia, Viola papilionacea*<br>● Raw GBIF records include the occurrence records searched by all the scientific names.<br>**Same state flower:**<br>● *Viola sororia* is the state flower of Illinois (IL), New Jersey (NJ), Rhode Island (RI), and Wisconsin (WI). |
| South Carolina (SC) | state flower | Yellow jessamine (*Gelsemium sempervirens*) | **Evidence of designating as the official state flower:**<br>● "...on the report of a select legislative commission consisting of Senator T. B. Butler of Gaffney, and Representatives G. B. Ellison, of Columbia, and T. S. Heyward, of Buffton, the General Assembly on February 1, 1924, adopted as the State flower the yellow jessamine, called also the Carolina jessamine [*Gelsemium sempervirens*]." [SC-1] |
| | state wildflower | Tall goldenrod (*Solidago altissima*) | **Evidence of designating as the official state wildflower:**<br>● "Goldenrod (*Solidago altissima*) is the official state wildflower." [SC-2] |
| South Dakota (SD) | state flower | America pasque flower (*Pulsatilla hirsutissima*) | **Evidence of designating as the official state flower:**<br>● From the South Dakota Codified Laws, Title 1, Chapter 1-6, Section 1-6-10. The floral emblem of this state shall be the American pasque flower (*Pulsatilla hirsutissima*) with the motto "I Lead." [SD-1]<br>**Synonym(s):**<br>● *Pulsatilla hirsutissima, Clematis hirsutissima*<br>● Raw GBIF records include the occurrence records searched by all the scientific names. |
| Tennessee | state flower | Purple Iris (*Iris versicolor*) | **Evidence of designating as the official state flower:** |

| | | | |
|---|---|---|---|
| (TN) | | | ● “In 1973 the 88th General Assembly, by Chapter 16, designated the passionflower the state wildflower and the iris the state cultivated flower.” [TN-1]<br>● “Legislation has never specified a particular variety of iris, but a variety of purple iris is usually depicted as representing the state's official cultivated flower.” [TN-1] |
| | state wildflower1 | Purple passionflower (*Passiflora incarnata*) | **Evidence of designating as the official state wildflower:**<br>● “The Passion Flower, genus Passiflora, is one of the official state wildflowers.” [TN-2] |
| | state wildflower2 | Tennessee purple coneflower (*Echinacea tennesseensis*) | **Evidence of designating as the official state wildflower:**<br>● “The Tennessee Coneflower, *Echinacea tennesseensis*, is one of the official state wildflowers.” [TN-2]<br>● “Tennessee recognized the exotic passion flower (*Passiflora incarnata*) as the official state flower in 1919 (chosen by the schoolchildren of Tennessee). But in 1933, Senate Joint Resolution 53 designated the iris as "the State Flower of Tennessee." In 1973 legislation was passed that distinguished the iris as the "state cultivated flower" and the passion flower as the "state wildflower." In 2012 a second wildflower was adopted as a symbol of Tennessee: the unique Tennessee coneflower. ” [TN-3] |
| Texas (TX) | state flower | Bluebonnet (*Lupinus* spp.) | **Evidence of designating as the official state flower:**<br>● “the Lupinus Texensis and any other variety of bluebonnet not heretofore recorded be recognized along with the *Lupinus subearnosus* as the official state flower of the State of Texas.” [TX-1]<br>**Multiple species:**<br>● According to Parsons et al. (Texas Cooperative Extension) publication: “Texas actually has five state flowers, more or less, and they are all bluebonnets.” [TX-2], including, *Lupinus subcarnosus*, *Lupinus texensis*, *Lupinus havardii*, *Lupinus concinnus*, *Lupinus plattensis*. So we modelled the potential distributions for the five species in Texas. |
| Utah (UT) | state flower | Sego lily (*Calochortus nuttallii*) | **Evidence of designating as the official state flower:**<br>● “The sego lily (*Calochortus nuttallii* Torr. & Gray) was approved as the official state flower of Utah by the Ninth Regular Session of the Legislature of the State of Utah. The legislation was signed by Governor William Spry on March 18, 1911.” [UT-1] |

| | | | |
|---|---|---|---|
| Vermont (VT) | state flower | Red clover (*Trifolium pratense*) | **Evidence of designating as the official state flower:**<br>● "On November 9, 1894, the red clover was adopted as the state flower by the Thirteenth Biennial Session of the General Assembly with an effective date of February 1, 1895." [VT-1] |
| Virginia (VA) | state flower | American dogwood (*Cornus florida*) | **Evidence of designating as the official state flower:**<br>● "The flower commonly known as the American Dogwood (*Cornus florida*), is hereby declared to be the floral emblem of the State of Virginia." [VA-1] |
| Washington (WA) | state flower | Coast rhododendron (*Rhododendron macrophyllum*) | **Evidence of designating as the official state flower:**<br>● "From the Washington Statutes, Title 1, Chapter 1.20, Section 1.20.030. The native species, *Rhododendron macrophyllum*, is hereby designated as the official flower of the state of Washington." [WA-1] |
| West Virginia (WV) | state flower | Rhododendron (*Rhododendron maximum*) | **Evidence of designating as the official state flower:**<br>● "A statewide referendum was finalized on November 26, 1902 when students expressed an overwhelming preference for the rhododendron. Big laurel won more than half (19,000) of the 36,000 votes cast posting solid backing in all West Virginia counties with the exception of Ohio County, where the goldenrod was favored. Following the landslide victory of big laurel, the flower was approved by the West Virginia Senate on January 23, 1903 and by the House on January 29, 1903 (House Joint Resolution No. 19)." [WV-1] |
| Wisconsin (WI) | state flower | Common blue violet (*Viola sororia*) | **Evidence of designating as the official state flower:**<br>● From the Wisconsin Statutes, Chapter 1, Section 1.10(3)(e). "The wood violet (*Viola papilionacea*) is the state flower." [WI-1]<br>**Synonym(s):**<br>● *Viola sororia, Viola papilionacea*<br>● Raw GBIF records include the occurrence records searched by all the scientific names.<br>**Same state flower:**<br>● Viola sororia is the state flower of Illinois (IL), New Jersey (NJ), Rhode Island (RI), and Wisconsin (WI). |
| Wyoming (WY) | state flower | Indian paintbrush (*Castilleja linariifolia*) | **Evidence of designating as the official state flower:**<br>● From the Wyoming Statutes, Title 8, Chapter 3, Section 8-3-104. "The *Castillija linariaefolia*, commonly called "the Indian paint brush," is the state flower of Wyoming." [WY-1] |
