## Supplement 2 for "Wilting Wildflowers and Bummed-Out Bees: Climate Change Threatens U.S. State Symbols"

**Supplement 2. Detailed documentation of the state insect species examined in this study.**

| State | Category | Common name (Scientific name) | Notes |
|---|---|---|---|
| Alabama (AL) | state agricultural insect | European honey bee (*Apis mellifera*) | **Evidence of designating as the official state agricultural insect:**<br>● "Queen honey bee designated as the Official State Agriculture Insect of Alabama." [AL-1]<br>**Same state insect:**<br>● *Apis mellifera* is the official state insect of 20 states, including Alabama (AL), Arkansas (AR), Georgia (GA), Kansas (KS), Kentucky (KY), Louisiana (LA), Maine (ME), Mississippi (MS), Missouri (MO), Nebraska (ME), New Jersey (NJ), North Carolina (NC), Oklahoma (OK), South Dakota (SD), Tennessee (TN), Texas (TX), Utah (UT), Vermont (VT), West Virginia (WV), Wisconsin (WI). |
| | state butterfly and mascot | Eastern tiger swallowtail (*Papilio glaucus*) | **Evidence of designating as the official state agricultural insect:**<br>● "The lovely eastern tiger swallowtail (*Papilio glaucas*) was designated the state butterfly and mascot of Alabama in 1989 at the request of Selma, Alabama's City Council (Selma is called "The Butterfly Capital of Alabama" and Selma's mascot is the tiger swallowtail butterfly). " [AL-2]<br>**Same state insect:**<br>● *Papilio glaucus* is the official state butterfly of 6 states, including Alabama (AL), Delaware (DE), Georgia (GA), North Carolina (NC), South Carolina (SC), Virginia (VA). |
| | state insect | Monarch butterfly (*Danaus plexippus*) | **Evidence of designating as the official state insect:**<br>● "Alabama designated the monarch butterfly (*Danaus plexippus*) as the official state insect in 1989". [AL-3]<br>**Same state insect:**<br>● *Danaus plexippus* is the official state insect/butterfly of 7 states, including Alabama (AL), Idaho (ID), Illinois (IL), Minnesota (MN), Texas (TX), Vermont (VT), West Virginia (WV). |
| Alaska (AK) | state insect | Four-spotted skimmer dragonfly (*Libellula quadrimaculata*) | **Evidence of designating as the official state insect:**<br>● "Alaska designated the four-spot skimmer dragonfly (*Libellula quadrimaculata*) as the official state insect in 1995." [AK-1] |

| | | | |
|---|---|---|---|
| Arizona (AZ) | state butterfly | Two-tailed swallowtail (*Papilio multicaudata*) | **Evidence of designating as the official state insect:**<br>● "Arizona designated the two-tailed swallowtail butterfly (*Papilio multicaudata*) as the official state butterfly in 2001 (also called two-tailed tiger swallowtail)." [AZ-1] |
| Arkansas (AR) | state butterfly | Diana fritillary butterfly (*Speyeria diana*) | **Evidence of designating as the official state butterfly:**<br>● "The Diana fritillary butterfly was designated the official butterfly of Arkansas in 2007." [AR-1] |
| | state insect | European honey bee (*Apis mellifera*) | **Evidence of designating as the official state insect:**<br>● "The honeybee was designated the official state insect of Arkansas in 1973." [AR-2]<br>**Same state insect:**<br>● *Apis mellifera* is the official state insect of 20 states, including Alabama (AL), Arkansas (AR), Georgia (GA), Kansas (KS), Kentucky (KY), Louisiana (LA), Maine (ME), Mississippi (MS), Missouri (MO), Nebraska (ME), New Jersey (NJ), North Carolina (NC), Oklahoma (OK), South Dakota (SD), Tennessee (TN), Texas (TX), Utah (UT), Vermont (VT), West Virginia (WV), Wisconsin (WI). |
| California (CA) | state butterfly | California dogface butterfly (*Zerene eurydice*) | **Evidence of designating as the official state insect:**<br>● "California designated the California dogface butterfly (*Zerene eurydice*) as the official state insect in 1972." [CA-1] |
| Colorado (CO) | state insect | Colorado hairstreak (*Hypaurotis crysalus*) | **Evidence of designating as the official state insect:**<br>● "The Colorado hairstreak butterfly (*Hypaurotis crysalus*) was designated the official state insect of Colorado in 1996 due to the steady lobbying of 4th graders from Wheeling Elementary in Aurora, Colorado (led by teacher Melinda Terry)." [CO-1] |
| Connecticut (CT) | state insect | European mantis (*Mantis religiosa*) | **Evidence of designating as the official state insect:**<br>● "The European praying mantis (*Mantis religiosa*) was designated the official state insect of Connecticut in 1977." [CT-1] |
| Delaware (DE) | state butterfly | Eastern tiger swallowtail (*Papilio glaucus*) | **Evidence of designating as the official state butterfly:**<br>● "The brilliant tiger swallowtail butterfly (*Pterourus glaucus*) was designated the official state butterfly of Delaware June 10,1999." [DE-1]<br>**Same state insect:**<br>● *Papilio glaucus* is the official state butterfly of 6 states, including Alabama (AL), Delaware (DE), Georgia (GA), North Carolina (NC), South Carolina (SC), Virginia (VA). |

| | | | |
|---|---|---|---|
| Florida (FL) | state butterfly | Zebra longwing (*Heliconius charithonia*) | **Evidence of designating as the official state insect:**<br>• "Florida designated the zebra longwing butterfly (*Heliconius charitonius*) as the official state butterfly in 1996." [FL-1] |
| Georgia (GA) | state insect | European honey bee (*Apis mellifera*) | **Evidence of designating as the official state insect:**<br>• "The honeybee was designated official state insect of Georgia in 1975 to acknowledge the insect's contribution to the state's economy through honey production and aiding pollination of more than 50 Georgia crops." [GA-1]<br>**Same state insect:**<br>• *Apis mellifera* is the official state insect of 20 states, including Alabama (AL), Arkansas (AR), Georgia (GA), Kansas (KS), Kentucky (KY), Louisiana (LA), Maine (ME), Mississippi (MS), Missouri (MO), Nebraska (ME), New Jersey (NJ), North Carolina (NC), Oklahoma (OK), South Dakota (SD), Tennessee (TN), Texas (TX), Utah (UT), Vermont (VT), West Virginia (WV), Wisconsin (WI). |
| | state butterfly | Eastern tiger swallowtail (*Papilio glaucus*) | **Evidence of designating as the official state butterfly:**<br>• "Georgia designated the tiger swallowtail (*Papilio glaucus*) as the official state butterfly in 1988." [GA-2]<br>**Same state insect:**<br>• *Papilio glaucus* is the official state butterfly of 6 states, including Alabama (AL), Delaware (DE), Georgia (GA), North Carolina (NC), South Carolina (SC), Virginia (VA). |
| Hawaii (HI) | state insect | Kamehameha butterfly (*Vanessa tameamea*) | **Evidence of designating as the official state insect:**<br>• "Hawaii designated pulelehua, also known as the kamehameha butterfly (*Vanessa tameamea*) as the official state insect in 2009." [HI-1] |
| Idaho (ID) | state insect | Monarch butterfly (*Danaus plexippus*) | **Evidence of designating as the official state insect:**<br>• "The Monarch Butterfly (*Danaus plexippus*) was adopted as the state insect by the state legislature in 1992." [ID-1]<br>**Same state insect:**<br>• *Danaus plexippus* is the official state insect/butterfly of 7 states, including Alabama (AL), Idaho (ID), Illinois (IL), Minnesota (MN), Texas (TX), Vermont (VT), West Virginia (WV). |
| Illinois (IL) | state insect | Monarch butterfly (*Danaus plexippus*) | **Evidence of designating as the official state insect:**<br>• "Illinois designated the iconic monarch butterfly as the official state insect in 1975, the result of lobbying by Illinois schoolchildren (a third grader from Decatur was the first to suggest the monarch as state insect)." [IL-1] |

| | | | |
|---|---|---|---|
| | | | **Same state insect:**<br>● *Danaus plexippus* is the official state insect/butterfly of 7 states, including Alabama (AL), Idaho (ID), Illinois (IL), Minnesota (MN), Texas (TX), Vermont (VT), West Virginia (WV). |
| Indiana (IN) | state insect | Say's firefly (*Pyractomena angulata*) | **Evidence of designating as the official state insect:**<br>● "Say's firefly (*Pyractomena angulata*) was designated the official state insect of Indiana on March 23, 2018, due to the perseverance of Cumberland Elementary School students in West Lafayette." [IN-1] |
| Kansas (KS) | state insect | European honey bee (*Apis mellifera*) | **Evidence of designating as the official state insect:**<br>● "Kansas designated the honeybee as official state insect in 1976 in response to a petition signed by over 2000 Kansas schoolchildren to make the honeybee the state insect." [KS-1]<br>**Same state insect:**<br>● *Apis mellifera* is the official state insect of 20 states, including Alabama (AL), Arkansas (AR), Georgia (GA), Kansas (KS), Kentucky (KY), Louisiana (LA), Maine (ME), Mississippi (MS), Missouri (MO), Nebraska (ME), New Jersey (NJ), North Carolina (NC), Oklahoma (OK), South Dakota (SD), Tennessee (TN), Texas (TX), Utah (UT), Vermont (VT), West Virginia (WV), Wisconsin (WI). |
| Kentucky (KY) | state agricultural insect | European honey bee (*Apis mellifera*) | **Evidence of designating as the official state insect:**<br>● "The honey bee (*Apis mellifera*) was designated the official state agricultural insect of Kentucky in 2010 (Kentucky also recognizes a state butterfly, adopted in 1990)." [KY-1]<br>**Same state insect:**<br>● *Apis mellifera* is the official state insect of 20 states, including Alabama (AL), Arkansas (AR), Georgia (GA), Kansas (KS), Kentucky (KY), Louisiana (LA), Maine (ME), Mississippi (MS), Missouri (MO), Nebraska (ME), New Jersey (NJ), North Carolina (NC), Oklahoma (OK), South Dakota (SD), Tennessee (TN), Texas (TX), Utah (UT), Vermont (VT), West Virginia (WV), Wisconsin (WI). |
| | state butterfly | Viceroy butterfly (*Limenitis archippus*) | **Evidence of designating as the official state butterfly:**<br>● "The viceroy butterfly (*Limenitis archippus*) was designated the official state butterfly of Kentucky in 1990" [KY-2] |

| | | | |
|---|---|---|---|
| Louisiana (LA) | state insect | European honey bee (*Apis mellifera*) | **Evidence of designating as the official state insect:**<br>● “Louisiana designated the honeybee (*Apis mallifera*) as official state insect in 1977.” [LA-1]<br>**Same state insect:**<br>● *Apis mellifera* is the official state insect of 20 states, including Alabama (AL), Arkansas (AR), Georgia (GA), Kansas (KS), Kentucky (KY), Louisiana (LA), Maine (ME), Mississippi (MS), Missouri (MO), Nebraska (ME), New Jersey (NJ), North Carolina (NC), Oklahoma (OK), South Dakota (SD), Tennessee (TN), Texas (TX), Utah (UT), Vermont (VT), West Virginia (WV), Wisconsin (WI). |
| Maine (ME) | state insect | European honey bee (*Apis mellifera*) | **Evidence of designating as the official state insect:**<br>● “The honeybee was designated the official state insect of Maine in 1975.” [ME-1]<br>**Same state insect:**<br>● *Apis mellifera* is the official state insect of 20 states, including Alabama (AL), Arkansas (AR), Georgia (GA), Kansas (KS), Kentucky (KY), Louisiana (LA), Maine (ME), Mississippi (MS), Missouri (MO), Nebraska (ME), New Jersey (NJ), North Carolina (NC), Oklahoma (OK), South Dakota (SD), Tennessee (TN), Texas (TX), Utah (UT), Vermont (VT), West Virginia (WV), Wisconsin (WI). |
| Maryland (MD) | state insect | Baltimore checkerspot butterfly (*Euphydryas phaeton*) | **Evidence of designating as the official state insect:**<br>● “Maryland designated the lovely Baltimore checkerspot butterfly (*Euphydryas phaeton*) as the official arthropodic emblem in 1973.” [MD-1] |
| Massachusetts (MA) | state insect or insect emblem | two-spotted Lady Beetle (*Adalia bipunctata)* | **Evidence of designating as the official state insect:**<br>● “The Ladybug, as official Insect of the Commonwealth of Massachusetts, came to fruition due to the efforts of a second grade class in Franklin.” [MA-1]<br>● “The most common Ladybug seen in Massachusetts is the two-spotted Lady Beetle (*Adalia bipunctata*). It's most easily recognized by the two distinct spots on its back. This Lady Beetle's head is black. Though Massachusetts law does not specify a scientific name, this is the accepted representative of the Ladybug family in Massachusetts.” [MA-1] |

| | | | |
|---|---|---|---|
| Michigan (MI) | state butterfly | Black swallowtail (*Papilio polyxenes*) | **Evidence of designating as the official state insect:**<br>● New bill introduced in 2023 to make this the state butterfly - hasn't been approved yet, but we included it in our study. [MI-1][MI-2]<br>**Same state insect:**<br>● *Papilio polyxenes* is the official state insect of 3 states, including Michigan(MI), New Jersey (NJ), Oklahoma (OK). |
| Minnesota (MN) | state bee | Rusty patched bumblebee (*Bombus affinis*) | **Evidence of designating as the official state bee:**<br>● “The rusty patched bumblebee, *Bombus affinis*, became Minnesota's state bee in 2019.” [MN-1] |
| | state butterfly | Monarch butterfly (*Danaus plexippus*) | **Evidence of designating as the official state insect:**<br>● “Minnesota adopted the lovely monarch butterfly (*Danaus plexippus*) as the official state butterfly in 2000.” [MN-1]<br>**Same state insect:**<br>● *Danaus plexippus* is the official state insect/butterfly of 7 states, including Alabama (AL), Idaho (ID), Illinois (IL), Minnesota (MN), Texas (TX), Vermont (VT), West Virginia (WV). |
| Mississippi (MS) | state butterfly | Spicebush swallowtail (*Papilio troilus*) | **Evidence of designating as the official state butterfly:**<br>● “Mississippi designated the spicebush swallowtail butterfly (*Papilio troilus*) as the official state butterfly in 1991.”[MS-1] |
| | state insect | European honey bee (*Apis mellifera*) | **Evidence of designating as the official state insect:**<br>● “The honeybee was designated the state Insect of Mississippi in 1980.” [MS-2]<br>**Same state insect:**<br>● *Apis mellifera* is the official state insect of 20 states, including Alabama (AL), Arkansas (AR), Georgia (GA), Kansas (KS), Kentucky (KY), Louisiana (LA), Maine (ME), Mississippi (MS), Missouri (MO), Nebraska (ME), New Jersey (NJ), North Carolina (NC), Oklahoma (OK), South Dakota (SD), Tennessee (TN), Texas (TX), Utah (UT), Vermont (VT), West Virginia (WV), Wisconsin (WI). |
| Missouri (MO) | state insect | European honey bee (*Apis mellifera*) | **Evidence of designating as the official state insect:**<br>● “Missouri designated the honeybee (*Apis mellifera*) as the official state insect in 1985.” [MO-1]<br>**Same state insect:**<br>● *Apis mellifera* is the official state insect of 20 states, including Alabama (AL), Arkansas (AR), Georgia (GA), Kansas (KS), Kentucky (KY), Louisiana (LA), Maine (ME), Mississippi (MS), Missouri (MO), Nebraska |

| | | | |
|---|---|---|---|
| | | | (ME), New Jersey (NJ), North Carolina (NC), Oklahoma (OK), South Dakota (SD), Tennessee (TN), Texas (TX), Utah (UT), Vermont (VT), West Virginia (WV), Wisconsin (WI). |
| Montana (MT) | state butterfly | Mourning cloak butterfly (*Nymphalis antiopa*) | **Evidence of designating as the official state insect:**<br>● "Montana designated the lovely mourning cloak butterfly (*Nymphalis antiopa*) as the official state butterfly in 2001." [MT-1] |
| Nebraska (NE) | state insect | European honey bee (*Apis mellifera*) | **Evidence of designating as the official state insect:**<br>● "The honeybee (*Apis mellifera*) was named the state insect by legislative action in 1975." [NE-1]<br>**Same state insect:**<br>● *Apis mellifera* is the official state insect of 20 states, including Alabama (AL), Arkansas (AR), Georgia (GA), Kansas (KS), Kentucky (KY), Louisiana (LA), Maine (ME), Mississippi (MS), Missouri (MO), Nebraska (ME), New Jersey (NJ), North Carolina (NC), Oklahoma (OK), South Dakota (SD), Tennessee (TN), Texas (TX), Utah (UT), Vermont (VT), West Virginia (WV), Wisconsin (WI). |
| Nevada (NV) | state insect | Vivid dancer damselfly (*Argia vivida*) | **Evidence of designating as the official state insect:**<br>● "Nevada designated the vivid dancer damselfly (*Argia vivida*) as the official state insect in 2009. [NV-1] |
| New Hampshire (NH) | state butterfly | Karner blue butterfly (*Plebejus melissa samuelis*) | **Evidence of designating as the official state butterfly:**<br>● "The endangered karner blue butterfly (*Lycaeides melissa samuelis*) was designated the official state butterfly of New Hampshire in 1992." [NH-1]<br>**Synonym(s):**<br>● *Lycaeides melissa samuelis*, *Plebejus melissa samuelis*, *Plebejus samuelis.*<br>● Raw GBIF records include the occurrence records searched by all the scientific names. |
| | state insect | Nine-spotted ladybug (*Coccinella novemnotata*) | **Evidence of designating as the official state insect:**<br>● "From the New Hampshire Revised Statutes, Title 1, Chapter 3, Section 3:11. The ladybug, also known as the ladybird and the lady beetle, is hereby designated as the official state insect of New Hampshire." [NH-2]<br>● The ladybug was never specified in legislation, but is listed as 9-spot here: https://www.netstate.com/states/symb/nh_symb.htm<br>**Same state insect:** |

| | | | |
|---|---|---|---|
| | | | • *Coccinella novemnotata* is the official state insect of 2 states, including New Hampshire (NH), New York (NY), |
| New Jersey (NJ) | state bug | European honey bee (*Apis mellifera*) | **Evidence of designating as the official state insect:**<br>• "The honeybee (*Apis mellifera*) was designated the official state bug of New Jersey in 1974 after encouragement given by a group of students from the Sunnybrae School in Hamilton Township." [NJ-1]<br>**Same state insect:**<br>• *Apis mellifera* is the official state insect of 20 states, including Alabama (AL), Arkansas (AR), Georgia (GA), Kansas (KS), Kentucky (KY), Louisiana (LA), Maine (ME), Mississippi (MS), Missouri (MO), Nebraska (ME), New Jersey (NJ), North Carolina (NC), Oklahoma (OK), South Dakota (SD), Tennessee (TN), Texas (TX), Utah (UT), Vermont (VT), West Virginia (WV), Wisconsin (WI). |
| | state butterfly | Black swallowtail (*Papilio polyxenes*) | **Same state insect:**<br>• *Papilio polyxenes* is the official state insect of 3 states, including Michigan(MI), New Jersey (NJ), Oklahoma (OK). |
| New Mexico (NM) | state butterfly | Sandia hairstreak (*Callophrys mcfarlandi*) | **Evidence of designating as the official state butterfly:**<br>• "The sandia hairstreak butterfly (*Callophrys mcfarlandi*) was designated the official state butterfly of New Mexico in 2003." [NM-1] |
| | state insect | Tarantula hawk wasp (*Pepsis grossa*) | **Evidence of designating as the official state insect:**<br>• "The Tarantula Hawk Wasp or Tarantula Hawk (*Pepsis formosa*) was selected because of an initiative from a classroom in Edgewood, NM. An elementary class and their teacher researched states which has selected state insects and then selected three insects for students around the state for which to vote. This species was then selected by the 39th legislature in 1989." [NM-2] |
| New York (NY) | state insect | Nine-Spotted Lady Beetle (*Coccinela novemnotata*) | **Evidence of designating as the official state insect:**<br>• "New York is the only state that specifies the scientific name in their designation of the Ladybug as the Official State Insect. They name the Nine-Spotted Lady Beetle (*Coccinela novemnotata*) as the Official State Insect." [NY-1]<br>• From the The Consolidated Laws of New York, STL - State, Article 6, Section 86. "The lady bug (*Coccinella novemnotata*) shall be the official insect of the state of New York." [NY-2] |

| | | | |
|---|---|---|---|
| North Carolina (NC) | state butterfly | Eastern tiger swallowtail (*Papilio glaucus*) | **Same state insect:**<br>● *Papilio glaucus* is the official state butterfly of 6 states, including Alabama (AL), Delaware (DE), Georgia (GA), North Carolina (NC), South Carolina (SC), Virginia (VA). |
| | state insect | European honey bee (*Apis mellifera*) | **Evidence of designating as the official state insect:**<br>● "North Carolina designated the European honeybee (*Apis mellifera*) as official state insect in 1973." [NC-1]<br>**Same state insect:**<br>● *Apis mellifera* is the official state insect of 20 states, including Alabama (AL), Arkansas (AR), Georgia (GA), Kansas (KS), Kentucky (KY), Louisiana (LA), Maine (ME), Mississippi (MS), Missouri (MO), Nebraska (ME), New Jersey (NJ), North Carolina (NC), Oklahoma (OK), South Dakota (SD), Tennessee (TN), Texas (TX), Utah (UT), Vermont (VT), West Virginia (WV), Wisconsin (WI). |
| North Dakota (ND) | state insect | Convergent lady beetle (*Hippodamia convergens*) | **Evidence of designating as the official state insect:**<br>● "North Dakota designated the convergent lady beetle (commonly called ladybug) as the official state insect in 2011." [ND-1]<br>**Same state insect:**<br>● *Hippodamia convergens* is the official state insect of 2 states, including North Dakota (ND) and Ohio (OH). |
| Ohio (OH) | state insect | Convergent lady beetle (*Hippodamia convergens*) | **Evidence of designating as the official state insect:**<br>● "The Ladybug was designated as the official state insect by Senate Concurrent Resolution 14, 111th General Assembly, 1975-1976 Session." [OH-1]<br>● In fact, the Convergent Lady Beetle is the official state insect of Ohio. [OH-2] |
| Oklahoma (OK) | state butterfly | Black swallowtail (*Papilio polyxenes*) | **Evidence of designating as the official state butterfly:**<br>● "Oklahoma designated the black swallowtail butterfly as the official state butterfly symbol in 1996."[OK-1]<br>**Same state insect:**<br>● *Papilio polyxenes* is the official state insect of 3 states, including Michigan(MI), New Jersey (NJ), Oklahoma (OK). |

| | | | |
|---|---|---|---|
| | state insect | European honey bee (*Apis mellifera*) | **Evidence of designating as the official state insect:**<br>● "Oklahoma designated the honeybee (*Apis mellifera*) as the official state insect in 1992." [OK-2]<br>**Same state insect:**<br>● *Apis mellifera* is the official state insect of 20 states, including Alabama (AL), Arkansas (AR), Georgia (GA), Kansas (KS), Kentucky (KY), Louisiana (LA), Maine (ME), Mississippi (MS), Missouri (MO), Nebraska (ME), New Jersey (NJ), North Carolina (NC), Oklahoma (OK), South Dakota (SD), Tennessee (TN), Texas (TX), Utah (UT), Vermont (VT), West Virginia (WV), Wisconsin (WI). |
| Oregon (OR) | state insect | Oregon swallowtail butterfly (*Papilio machaon oregonia*) | **Evidence of designating as the official state insect:**<br>● "The Oregon swallowtail butterfly was adopted as the Oregon's official insect with the approval of Senate Concurrent Resolution No. 6 by the Oregon Legislature." [OR-1] |
| Pennsylvania (PA) | state insect | Pennsylvania firefly (*Photuris pensylvanica*) | **Evidence of designating as the official state insect:**<br>● From the The Pennsylvania Statutes, Title 71, Chapter 6, Section 1010. "The firefly (Lampyridae Coleoptera) of the species *Photuris pensylvanica* De Geer is hereby selected, designated and adopted as the official insect of the Commonwealth of Pennsylvania." [PA-1] |
| Rhode Island (RI) | state insect | American burying beetle (*Nicrophorus americanus*) | **Evidence of designating as the official state insect:**<br>● "Rhode Island designated the endangered American burying beetle as the official state insect on July 14, 2015 thanks to the third graders at St. Michael's Country Day School in Newport." [RI-1] |
| South Carolina (SC) | state butterfly | Eastern tiger swallowtail (*Papilio glaucus*) | **Evidence of designating as the official state butterfly:**<br>● "South Carolina designated the tiger swallowtail (*Pterourus glaucus*) as the official state butterfly in 1994." [SC-1]<br>**Same state insect:**<br>● *Papilio glaucus* is the official state butterfly of 6 states, including Alabama (AL), Delaware (DE), Georgia (GA), North Carolina (NC), South Carolina (SC), Virginia (VA). |
| | state insect | Carolina mantis (*Stagmomantis carolina*) | **Evidence of designating as the official state insect:** |

| | | | |
|---|---|---|---|
| | | | • From the The South Carolina Code of Laws, Title 1, Chapter , Article 9, section 1-1-645. “The Carolina mantid, Stagmomantis carolina (Johannson) , or praying mantis, is the official insect of the State.” [SC-2] |
| South Dakota (SD) | state insect | European honey bee (*Apis mellifera*) | **Evidence of designating as the official state insect:**<br>• “South Dakota, a leader in honey production, designated the honeybee as official state insect in 1978.” [SD-1]<br>**Same state insect:**<br>• *Apis mellifera* is the official state insect of 20 states, including Alabama (AL), Arkansas (AR), Georgia (GA), Kansas (KS), Kentucky (KY), Louisiana (LA), Maine (ME), Mississippi (MS), Missouri (MO), Nebraska (ME), New Jersey (NJ), North Carolina (NC), Oklahoma (OK), South Dakota (SD), Tennessee (TN), Texas (TX), Utah (UT), Vermont (VT), West Virginia (WV), Wisconsin (WI). |
| Tennessee (TN) | state agricultural insect | European honey bee (*Apis mellifera*) | **Evidence of designating as the official state agricultural insect:**<br>• “The honeybee was recognized as the official state agricultural insect of Tennessee in 1990.” [TN-1]<br>**Same state insect:**<br>• *Apis mellifera* is the official state insect of 20 states, including Alabama (AL), Arkansas (AR), Georgia (GA), Kansas (KS), Kentucky (KY), Louisiana (LA), Maine (ME), Mississippi (MS), Missouri (MO), Nebraska (ME), New Jersey (NJ), North Carolina (NC), Oklahoma (OK), South Dakota (SD), Tennessee (TN), Texas (TX), Utah (UT), Vermont (VT), West Virginia (WV), Wisconsin (WI). |
| | state butterfly | Zebra swallowtail (*Eurytides marcellus*) | **Evidence of designating as the official state butterfly:**<br>• “The strikingly beautiful zebra swallowtail butterfly (*Euryotides marcellus*) was designated the official state butterfly of Tennessee in 1995.” [TN-2] |
| | state insect1 | 7-spotted ladybug (*Coccinella septempunctata*) | **Evidence of designating as the official state insect:**<br>• “The official state insects were designated by Public Chapter 292 of the Acts of 1975. They are the well-known firefly, or lightning bug beetle, and the ladybeetle, more commonly known as the ladybug or ladybird beetle.” [TN-3]<br>• “The ladybeetle, more commonly called ladybug or ladybird beetle, is the popular name given the Coccinella 7.” [TN-3] |
| | state insect2 | Common eastern firefly (*Photinus pyralis*) | **Evidence of designating as the official state insect:**<br>• From the Tennessee Code Annotates, Title 4, Chapter 1, Part 3, Section 4-1-308. “The well-known firefly, or lightning bug beetle, and the ladybird |

| | | | |
|---|---|---|---|
| | | | beetle, commonly known as the ladybug, are hereby designated as the official state insects." [TN-4]<br>● "The most familiar species of firefly in Tennessee is *Photinus pyralls*." [TN-5] |
| Texas (TX) | state insect | Monarch butterfly (*Danaus plexippus*) | **Evidence of designating as the official state insect:**<br>● "Texas designated the monarch butterfly (*Danaus plexippus*) as the official state insect in 1995." [TX-1]<br>**Same state insect:**<br>● *Danaus plexippus* is the official state insect/butterfly of 7 states, including Alabama (AL), Idaho (ID), Illinois (IL), Minnesota (MN), Texas (TX), Vermont (VT), West Virginia (WV). |
|  | state pollinator | European honey bee (*Apis mellifera*) | **Evidence of designating as the official state pollinator:**<br>● "Texas designated the western honey bee (*Apis mellifera*) as the official state pollinator in 2015." [TX-2]<br>**Same state insect:**<br>● *Apis mellifera* is the official state insect of 20 states, including Alabama (AL), Arkansas (AR), Georgia (GA), Kansas (KS), Kentucky (KY), Louisiana (LA), Maine (ME), Mississippi (MS), Missouri (MO), Nebraska (ME), New Jersey (NJ), North Carolina (NC), Oklahoma (OK), South Dakota (SD), Tennessee (TN), Texas (TX), Utah (UT), Vermont (VT), West Virginia (WV), Wisconsin (WI). |
| Utah (UT) | state insect | European honey bee (*Apis mellifera*) | **Evidence of designating as the official state insect:**<br>● "Utah designated the honeybee as official state insect in 1983 due to the lobbying efforts of a fifth grade class." [UT-1]<br>**Same state insect:**<br>● *Apis mellifera* is the official state insect of 20 states, including Alabama (AL), Arkansas (AR), Georgia (GA), Kansas (KS), Kentucky (KY), Louisiana (LA), Maine (ME), Mississippi (MS), Missouri (MO), Nebraska (ME), New Jersey (NJ), North Carolina (NC), Oklahoma (OK), South Dakota (SD), Tennessee (TN), Texas (TX), Utah (UT), Vermont (VT), West Virginia (WV), Wisconsin (WI). |
| Vermont (VT) | state butterfly | Monarch butterfly (*Danaus plexippus*) | **Evidence of designating as the official state butterfly:**<br>● "Vermont designated the monarch (*Danaus plexippus*) as the official state butterfly in 1987". [VT-1]<br>**Same state insect:** |

| | | | |
|---|---|---|---|
| | | | • *Danaus plexippus* is the official state insect/butterfly of 7 states, including Alabama (AL), Idaho (ID), Illinois (IL), Minnesota (MN), Texas (TX), Vermont (VT), West Virginia (WV). |
| | state insect | European honey bee (*Apis mellifera*) | **Evidence of designating as the official state insect:**<br>• "Vermont designated the honeybee as official state insect in 1978." [VT-2]<br>**Same state insect:**<br>• *Apis mellifera* is the official state insect of 20 states, including Alabama (AL), Arkansas (AR), Georgia (GA), Kansas (KS), Kentucky (KY), Louisiana (LA), Maine (ME), Mississippi (MS), Missouri (MO), Nebraska (ME), New Jersey (NJ), North Carolina (NC), Oklahoma (OK), South Dakota (SD), Tennessee (TN), Texas (TX), Utah (UT), Vermont (VT), West Virginia (WV), Wisconsin (WI). |
| Virginia (VA) | state insect | Tiger swallowtail butterfly (*Papilio glaucus*) | **Evidence of designating as the official state insect:**<br>• "Virginia designated the tiger swallowtail butterfly (*Papilio glaucus* Linnaeus) as the official state insect in 1991." [VA-1]<br>**Same state insect:**<br>• *Papilio glaucus* is the official state butterfly of 6 states, including Alabama (AL), Delaware (DE), Georgia (GA), North Carolina (NC), South Carolina (SC), Virginia (VA). |
| Washington (WA) | state insect | Green darner dragonfly (*Anax junius*) | **Evidence of designating as the official state insect:**<br>• "The common green darner dragonfly, *Anax junius* Drury, is hereby designated as the official insect of the state of Washington." [WA-1] |
| West Virginia (WV) | state butterfly | Monarch butterfly (*Danaus plexippus*) | **Evidence of designating as the official state butterfly:**<br>• "West Virginia designated the monarch butterfly (*Danaus plexippus*) as the official state butterfly in 1995 (seven states have recognized the monarch butterfly as a state symbol)." [WV-1]<br>**Same state insect:**<br>• *Danaus plexippus* is the official state insect/butterfly of 7 states, including Alabama (AL), Idaho (ID), Illinois (IL), Minnesota (MN), Texas (TX), Vermont (VT), West Virginia (WV). |
| | state insect | European honey bee (*Apis mellifera*) | **Evidence of designating as the official state insect:**<br>• "In 2002, after legislators struggled between the lady beetle and the honey bee, West Virginia finally designated the honeybee (*Apis mellifera*) as the official state insect." [WV-2]<br>**Same state insect:** |

| | | | |
|---|---|---|---|
| | | | ● *Apis mellifera* is the official state insect of 20 states, including Alabama (AL), Arkansas (AR), Georgia (GA), Kansas (KS), Kentucky (KY), Louisiana (LA), Maine (ME), Mississippi (MS), Missouri (MO), Nebraska (ME), New Jersey (NJ), North Carolina (NC), Oklahoma (OK), South Dakota (SD), Tennessee (TN), Texas (TX), Utah (UT), Vermont (VT), West Virginia (WV), Wisconsin (WI). |
| Wisconsin (WI) | state insect | European honey bee (*Apis mellifera*) | **Evidence of designating as the official state insect:**<br>● "Wisconsin designated the honeybee as official state insect in 1977." [WI-1]<br>**Same state insect:**<br>● *Apis mellifera* is the official state insect of 20 states, including Alabama (AL), Arkansas (AR), Georgia (GA), Kansas (KS), Kentucky (KY), Louisiana (LA), Maine (ME), Mississippi (MS), Missouri (MO), Nebraska (ME), New Jersey (NJ), North Carolina (NC), Oklahoma (OK), South Dakota (SD), Tennessee (TN), Texas (TX), Utah (UT), Vermont (VT), West Virginia (WV), Wisconsin (WI). |
| Wyoming (WY) | state insect | Sheridan's green hairstreak (*Callophrys sheridanii*) | **Evidence of designating as the official state insect:**<br>● "*Callophrys sheridanii*, commonly known as Sheridan's green hairstreak butterfly, is the state butterfly of Wyoming." [WY-1] |
