## Supplementary Figuer S1-S3 for "Wilting Wildflowers and Bummed-Out Bees: Climate Change Threatens U.S. State Symbols"

**Figure S1. State flowers:** Geographical distribution of different types of climate change effects on the habitat suitability (HSVs) for state flowers. We selected five representative state flowers to illustrate the geographical distribution of habitat suitability under historical (1981-2010) and future (2071-2100, SSP1-2.6 and SSP5-8.5) climate conditions. The five state flowers are selected based on the predictions under SSP5-8.5 scenario. Additionally, we plotted the geographical distribution and probability density of $\Delta$HSV to highlight the changes in habitat suitability due to climate change.

**Figure S2. State insects:** Geographical distribution of different types of climate change effects on the habitat suitability (HSVs) for state insects. We selected six representative state insects to illustrate the geographical distribution of habitat suitability under historical (1981-2010) and future (2071-2100, SSP1-2.6 and SSP5-8.5) climate conditions. The six state insects are selected based on the predictions under SSP5-8.5 scenario. Additionally, we plotted the geographical distribution and probability density of $\Delta$HSV to highlight the changes in habitat suitability due to climate change.

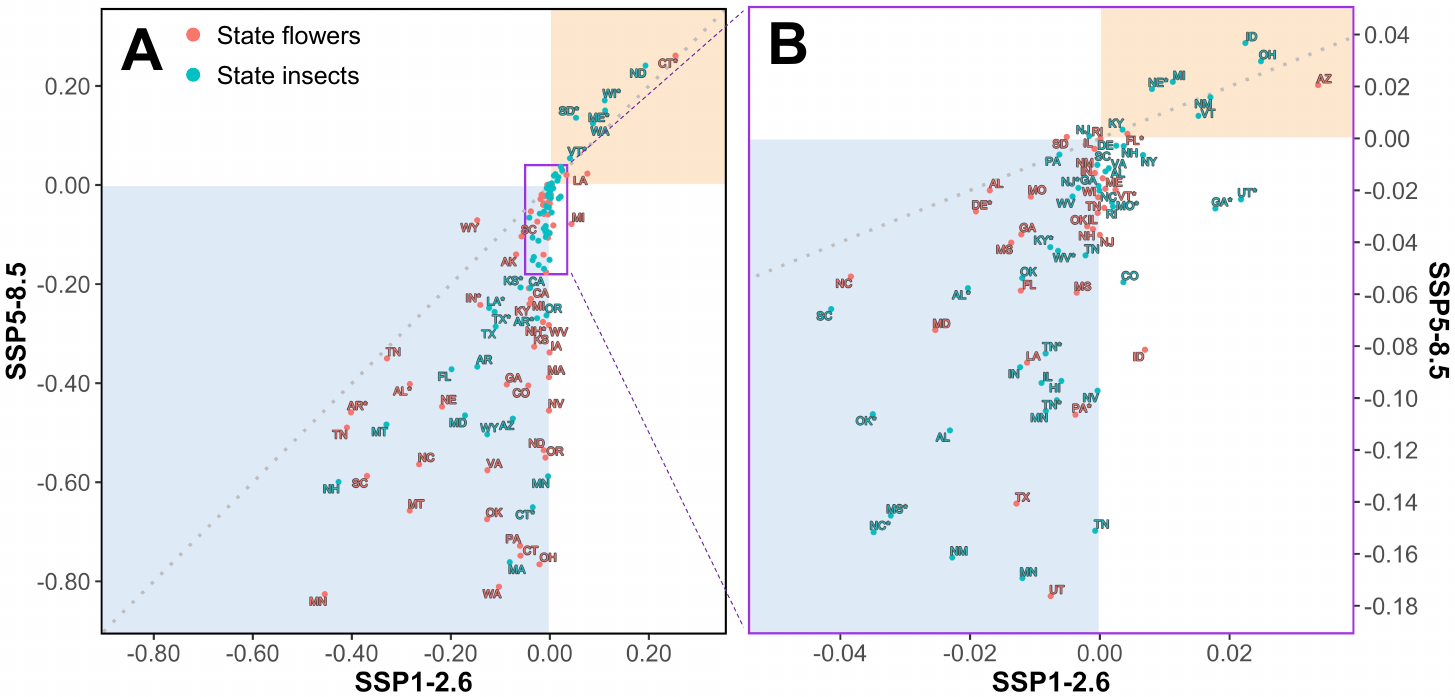

**Figure S3.** Changes in habitat suitability for state flowers and insects under climate change. The location of each text box in the plot indicates the differences in the median habitat suitability value between future and historical conditions for each state species in its designated state, under the SSP1-2.6 and SSP5-8.5 scenarios. Non-native species are indicated by red * in y-axis labels.
